# Characterization of the substitution hotspots in SARS-CoV-2 genome using BioAider and detection of a SR-rich region in N protein providing further evidence of its animal origin

**DOI:** 10.1101/2020.06.04.135293

**Authors:** Zhi-Jian Zhou, Ye Qiu, Xing-Yi Ge

## Abstract

The novel human coronavirus (SARS-CoV-2) causes the coronavirus disease 2019 (COVID-19) pandemic worldwide. The increasing sequencing data have shown abundant single nucleotide variations in SARS-CoV-2 genome. However, it is difficult to quickly analyze genomic variation and screen key mutations of SARS-CoV-2. In this study, we developed a visual program, named BioAider, for quick and convenient sequence annotation and mutation analysis on multiple genome-sequencing data. Using BioAider, we conducted a comprehensive genome variation analysis on 3,240 sequences of SARS-CoV-2 genome. Herein, we detected 14 substitution hotspots within SARS-CoV-2 genome, including 10 non-synonymous and 4 synonymous ones. Among these hotspots, NSP13-Y541C was predicted to be a crucial substitution which might affect the unwinding activity of NSP13, a key protein for viral replication. Besides, we also found 3 groups of potentially linked substitution hotspots which were worth further study. In particular, we discovered a SR-rich region (aa 184-204) on the N protein of SARS-CoV-2 distinct from SARS-CoV, indicating more complex replication mechanism and unique N-M interaction of SARS-CoV-2. Interestingly, the quantity of SRXX repeat fragments in the SR-rich region well reflected the evolutionary relationship among SARS-CoV-2 and SARS-CoV-2 related animal coronaviruses, providing further evidence of its animal origin. Overall, we developed an efficient tool for rapid identification of mutations, identified substitution hotspots in SARS-CoV-2 genomes, and detected a distinctive polymorphism SR-rich region in N protein. This tool and the detected hotspots could facilitate the viral genomic study and may contribute for screening antiviral target sites.

## Introduction

The severe pandemic of Coronavirus Disease 2019 (COVID-19) is caused by a 2019 novel coronavirus (2019-nCoV) which was first detected and characterized in Wuhan, China at the end of 2019 [1, 2]. Later, the name of this virus was suggested as severe acute respiratory syndrome coronavirus 2 (SARS-CoV-2) by the International Committee on Taxonomy of Viruses (ICTV) [3]. SARS-CoV-2 is a new pathogen and showing high infectiousness, fast spread, partial asymptomatic infection, and other new features [4, 5]. At present, the virus has been detected in the most countries and regions around the world, threatening public health, normal social life, and economy. SARS-CoV-2 has infected millions of people with more than 250,000 deaths worldwide as of May 8th, 2020 (https://www.who.int/emergencies/diseases/novel-coronavirus-2019/situation-reports/). Most patients infected with SARS-CoV-2 mainly present lower respiratory tract infections, accompanied by fever, dry cough, dyspnea [6]. So far, there is no specific drug to treat SARS-CoV-2 infection. Moreover, the vaccine against SARS-CoV-2 is still on the way. The world will be still facing COVID-19 for a long time.

Coronaviruses are enveloped virus with a non-segmented positive sense RNA genome, the full-length genome is the largest among RNA viruses [7]. The first ORF encodes a polyprotein 1ab (pp1ab, ORF1ab) and approximately occupied the first two-thirds of the genome, the remainder are structural and non-structural proteins [8]. The pp1ab usually is hydrolyzed into 16 non-structural proteins (NSP1-16) by one or two papain-like protease (PLPs) in NSP3 and 3C-like protease (3CL^pro^) in NSP5, to promote viral RNA replication and transcription [9, 10]. For these non-structural proteins in pp1ab, NSP2 can inhibit two host proteins of PHB1 and PHB2, which play an important role in viral infection [11]. The PLPs of NSP3 hydrolyze the N-terminal of pp1ab to NSP1 ~ NSP4, 3CL^pro^ of NSP5 binds 11 conserved Q-S dipeptide sites in pp1ab to produce 12 mature non-structural proteins (NSP5 ~ NSP16) [9, 12]. NSP12 is a key component, also known as the RNA-dependent RNA polymerase (RdRp) and is crucial in the replication and transcription cycle of coronaviruses [13]. Currently, RdRp is considered to be the main target of antiviral drugs for SARS-CoV-2[14]. NSP13 owns NTPase/Helicase activity and can unwind double-stranded RNA and DNA helix [15]. Due to the conservation of NSP13 sequence and necessity in all coronaviruses species, it is considered an ideal target for the development of antiviral drugs for SARS-CoV [16, 17]. The main four structural proteins of coronaviruses are the spike protein (S), small envelope protein (E), membrane protein (M) and nucleocapsid protein (N) [18]. S protein interacts with molecular receptors to mediate cell membrane fusion, allowing viruses to enter host cells [19]. E and M proteins are involved in the assembly of the virus, which is related to the formation and release of the viral envelope, N protein plays an important role in virus replication and pathogenesis [19, 20].

Based on seven conserved replicase domains in pp1ab (polyprotein-1ab) for the classification of coronavirus, SARS-CoV-2 belongs to the *Sarbecovirus* subgenus of the genus *Betacoronavirus* in the subfamily *Orthocoronavirinae* together with human SARS-CoV and bat various SARSr-CoVs (SARS-related CoVs), and SARS-CoV-2 is highly similar to bat coronavirus Bat-CoV-RaTG13 genetically [1, 21, 22]. Similar to SARS-CoV, SARS-CoV-2 uses the angiotensin converting enzyme II (ACE2) as receptor, besides, the serine protease TMPRSS2 plays an important role in activating spike (S) protein [23]. Some novel features of SARS-CoV-2 has been revealed, like a furin protease cleavage site in the Spike which may be related to viral transmissibility [24]. Recently, a novel bat-derived coronavirus RmYN02 with the similar furin insertion in the S was found which emphasized the bat origin of the virus [25].

As a member of RNA virus, the RNA-dependent RNA polymerase (RdRp) encoded by coronavirus lacks the proofreading capability, which leading to high mis-incorporation rate during replication even with the help of exoribonuclease like ExoN [26]. Together with the genome recombination events, it could cause viral diversity and promote the transmission and disease of coronavirus [27]. At the same time, in order to better adapt to the host, the virus usually mutates continuously under the selection pressure of the host. In particular, some non-synonymous substitution sites with high frequency are often experience strong positive selection [28]. Considering the continuous increase of infected people and the high variability of the virus, it is important to pay attention to the genomic changes of SARS-CoV-2. Recently, 7 substitution hotspots in SARS-CoV-2, ORF1ab-G10818T (ORF1ab-L3606F), ORF1ab-C8517T, ORF3a-G752T (ORF3a-G251V) S-A1841G (D614G), G171T (Q57H), ORF8-T251C (ORF8-L84S) and N-GGG608_609_610AAC (**N**-RG203_204KR) have been reported [29–32]. In addition, the non-synonymous substitution of ORF3a-G251V and ORF8-L84S both cause the amino acid (aa) polarity changing, which may affect the conformation of the protein and lead to function altering [31].

Despite these recent discoveries, in order to deal with the continuous variation of SARS-CoV-2 and the accumulation of sequencing data, methods for quick and efficient analysis of the mutations feature and substitutions hotspots of SARS-CoV-2 genome are urgently required. However, analyzing a large number of variant sequences is difficult for biological or clinical expert without bioinformatics and programming skills. Some tools such as MutPred can analyze variations by entering amino acid sequence, but unable to handle synonymous substitutions and consecutive nucleotide mutations like dinucleotide and trinucleotide substitution [33]. In this respect, we have developed a visualization analysis tool, Bioinformatics Aider V1.0 (BioAider V1.0, https://github.com/ZhijianZhou01/BioAider/releases). BioAider showed high efficiency and convenient in gene annotation and mutation screening on multiple genome-sequencing data.

In this research, we collected 3240 complete genome sequences of SARS-CoV-2 from 64 different countries and regions, sampling time from December 24, 2019 to April 1, 2020. We conducted a detailed genome mutation analysis using BioAider. We identified 14 substitution hotspots and 3 groups of possible linkage substitutions. Especially, we found distinctive polymorphism on SR-rich region of N protein in SARS-CoV-2 and related coronaviruses in other animals. Our work provides a new tool for recognition of the variation and evolution of SARS-CoV-2, contributes to research the replication and pathogenic mechanism of SARS-CoV-2, and gives further evidence for the animal origin of SARS-CoV-2.

## Results

### BioAider

For these mutation analysis results of Bioaider V1.0, the two main results are the summary of mutation sites (Fig. 1A) and the detailed log files (Fig. 1B) for positioning mutated strain. BioAider puts complex algorithms and fault-tolerant processing mechanisms inside, presenting users a very simple interface and prompts are added to interface controls, which is extremely friendly for users. Especially, we fully considered the time complexity and space complexity and optimization related algorithms, which makes BioAider owns higher processing efficiency. More importantly, it will continue to be in development, we will update the version for Linux systems and add new features later. As a demonstration, here we used BioAider for the mutations analysis of viral genome sequences.

**Fig. 1.**
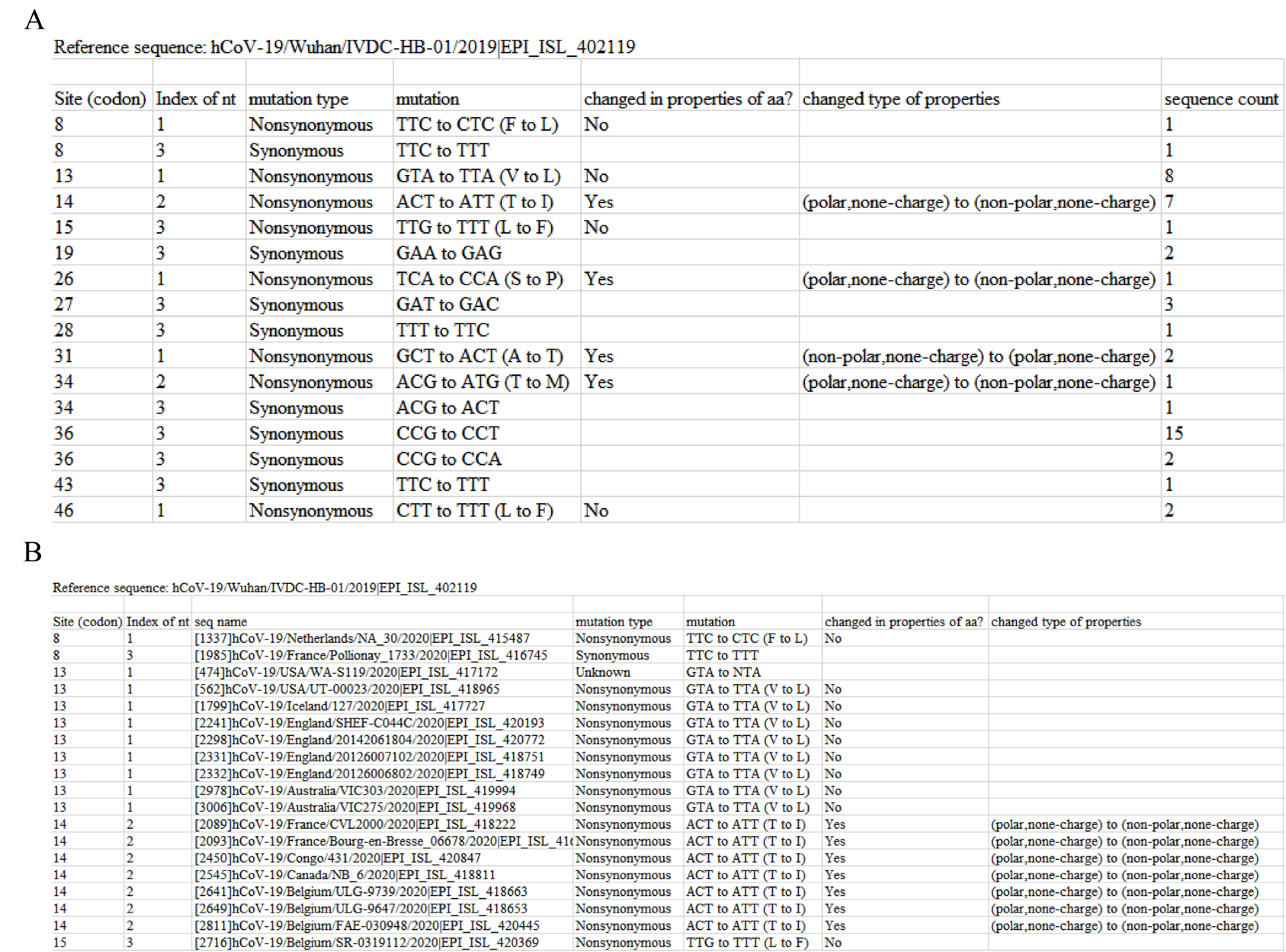
Examples of sequence analysis results by BioAider. (A) Summary of mutation site information. (B). Detailed log files for query.

### Mutation sites in SARS-CoV-2 genomes

In this study, ‘substitution’ refers to synonymous or non-synonymous substitution, and ‘mutation’ includes all the genomic variance. The frequency of mutation (or substitution) refers to the number of mutant strains compared to the reference strain sequence in this study.

Compared with the early viral sampling strains (EPI_ISL_402119) in GISAID database, a total of 2152 mutation sites (regardless of insertions or deletions) were identified in 3239 SARS-CoV-2 sequenced whole genomes, accounting for 7.36% of SARS-CoV-2 full-length gene (Table 1). The number of synonymous and non-synonymous substitution sites was counted as 784 (2.68%) and 1335 (4.57%), respectively. Among these non-synonymous substitution sites, 738 resulted in changes in amino acid properties. Besides, 12 sites contained both synonymous and non-synonymous substitution depending on the strains, and a total of 21 termination mutation sites were also detected (Table 1). In all the coding genes, three with the of the most mutation sites were ORF1ab, spike (S) and nucleocapsid (N) gene, the number of mutations sites were 1400, 296 and 168, respectively, accounting for 6.58%, 7.75% and 13.36% in their corresponding gene length (Table 1). The gene with the highest proportion of mutation sites was ORF10, containing 16 mutation sites were in 114 bases which occupied 14.04% of ORF10. All the mutation sites and viral starins of SARS-CoV-2 detected by BioAider in this study were summarized in sTable 1 and sTable 2.

**Table 1.** Mutation sites of SARS-CoV-2 based on 3239 sequenced strains

| Genes |  | Length | Mutation sites | S(p) / N(p) / S-N(p) / Termination(p) |
| --- | --- | --- | --- | --- |
|  |  |  | Count/Proportion |  |
| ORF1ab | NSP1 | 540 | 48/8.88% | 24(4.44%)/24(4.44%)/0/0 |
|  | NSP2 | 1914 | 168/8.77% | 46(2.40%)/122(6.37%)/0/0 |
|  | NSP3 | 5835 | 384/6.58% | 130(2.23%)/253 (4.33%)/1(0.02%)/0 |
|  | NSP4 | 1500 | 99/6.60% | 46(3.07%)/53(3.53%)/0/0 |
|  | NSP5 | 918 | 42/4.58% | 19(2.07%)/23(2.51%)/0/0 |
|  | NSP6 | 870 | 71/8.16% | 30(3.45%)/41(4.71%)/0/0 |
|  | NSP7 | 249 | 19/7.63% | 9(3.61%)/10(4.02%)/0/0 |
|  | NSP8 | 594 | 27/4.55% | 9(1.52%)/16(2.69%)/1(0.17%)/1(0.17%) |
|  | NSP9 | 339 | 21/6.19% | 11(3.24%)/10(2.95%)/0/0 |
|  | NSP10 | 417 | 27/6.48% | 10(2.40%)/14(3.36%)/1(0.24%)/2(0.48%) |
|  | NSP12 | 2796 | 157/5.61% | 58(2.07%)/96(3.43%)/0/3(0.11%) |
|  | NSP13 | 1803 | 109/6.05% | 42(2.33%)/67(3.72%)/0/0 |
|  | NSP14 | 1581 | 99/6.25% | 41(2.59%)/56(3.54%)/1(0.06%)/1(0.06%) |
|  | NSP15 | 1038 | 62/5.97% | 18(1.73%)/43(4.14%)/1(0.10%)/0 |
|  | NSP16 | 894 | 67/7.49% | 27(3.02%)/39(4.36%)/0/1(0.11%) |
|  | **Count** | 21288 | 1400/6.57% | 520(2.44%)/867(4.07%)/5(0.02%)/8(0.04%) |
| S |  | 3819 | 296/7.75% | 105(2.75%)/186(4.87%)/3(0.08%)/2(0.05%) |
| ORF3a |  | 825 | 94/11.39% | 31(3.75%)/63(7.64%)/0/0 |
| E |  | 225 | 22/9.78% | 9(4.00%)/13(5.78%)/0/0 |
| M |  | 666 | 44/6.61% | 28(4.21%)/16(2.40%)/0/0 |
| ORF6 |  | 183 | 22/12.01% | 4(2.18%)/16(8.74%)/0/2(1.09%) |
| ORF7a |  | 363 | 40/11.02% | 12(3.31%)/22(6.06%)/1(0.27%)/5(1.38%) |
| ORF7b |  | 129 | 11/8.52% | 3(2.33%)/7(5.42%)/0/1(0.77%) |
| ORF8 |  | 363 | 39/10.75% | 10(2.75%)/26(7.16%)/0/3(0.83%) |
| N |  | 1257 | 168/13.36% | 58(4.61%)/107(8.51%)/3(0.24%)/0 |
| ORF10 |  | 114 | 16/14.04% | 4(3.51%)/12(10.53%)/0/0 |
| Complete |  | 29232 | 2152/7.36% | 784(2.68%)/1335(4.57%)/12(0.04%)/21(0.07%) |

### Substitution frequency distribution of synonymous or nonsynonymous sites

To assess the overall substitution frequency of these mutated sites, we divided the substitution frequency into six different groups, and drew the frequency spectra of 2119 substitution sites (784 synonymous and 1335 nonsynonymous) of 3239 sequenced strains (Fig. 2). The result shows that the substitution sites of non-synonymous were always over synonymous except for the fifth group. Besides, a large number of sites with substitution frequencies between 1 and 5, and more than half of the substitution sites only are observed in a single strain. We also found 60 sites owning a substitution frequency greater than 20, including 40 non-synonymous substitution sites and 20 synonymous substitution sites. The substitution frequency distribution of synonymous or nonsynonymous sites for each codon gene are in sFigure 1.

**Fig. 2.**
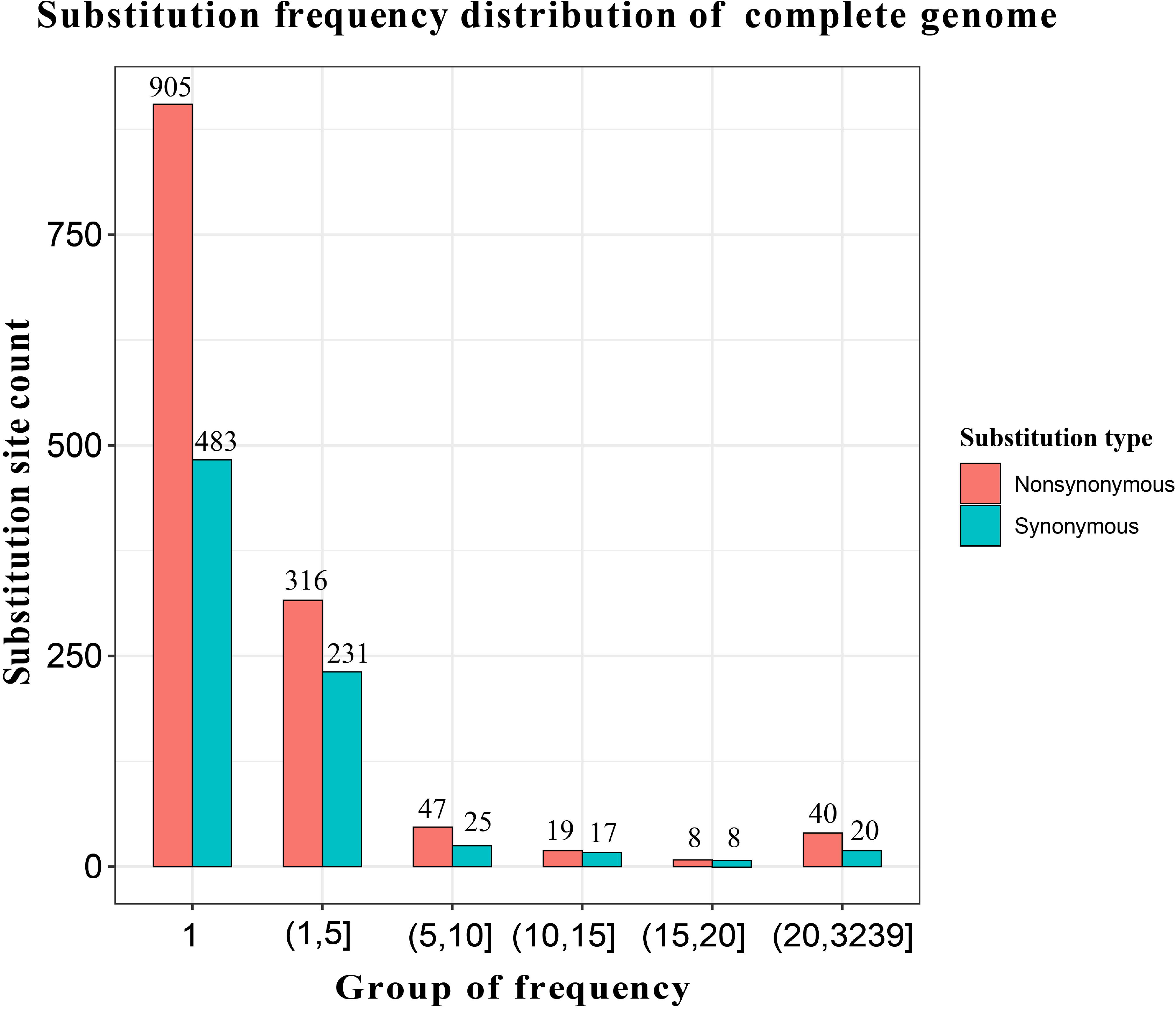
Substitution frequency of synonymous or nonsynonymous sites in SARS-CoV-2 genome. The frequency of X axis indicates the number of variant strains at the same substitution site, and the number one of first group represents that only one strain was mutated at this site. The Y axis represents the count of substitution sites corresponding to the range of frequency. The frequency spectra was drawn using BioAider by specifying five different groups.

### Substitution hotspots in SARS-CoV-2 genomes

Totally, 2119 substitution sites were detected in 3239 sequenced strains, most of them with a lower frequency. We defined the site with substitution frequency over 200 as the substitution hotspot, thus far, 14 substitution hotspots were identified which distributed in ORF1ab, S, ORF3a, ORF8 and N gene (Fig. 3). Among these substitution hotspots, 10 sites were non-synonymous and 4 were synonymous. As for these non-synonymous substitution hotspots, 6 substitution sites caused a change on polarity or chargeability of amino acid. Especially, there was a trinucleotide substitution hotspot on the N gene and caused two amino acid changes (Fig. 3).

**Fig. 3.**
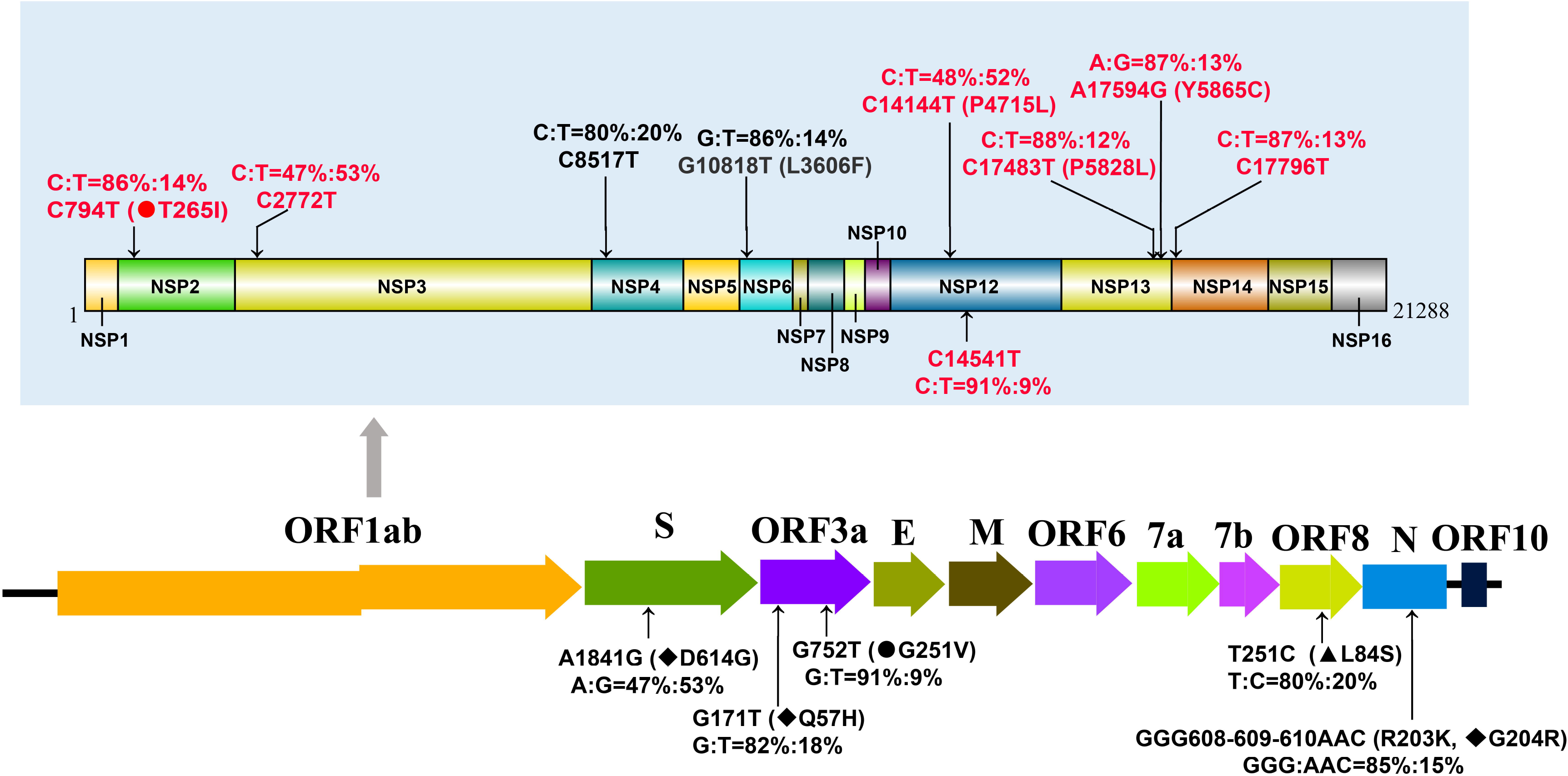
Substitution hotspots in 3239 SARS-CoV-2 full genomic sequences. The value of the substitution site indicates the position on the corresponding gene, and showed the two main bases and ratio at substitution hotspots of 3239 sequenced strains. In particular, the substitution hotspots in N gene was a trinucleotide mutation. Round represented polar aa changed to non-polar aa, triangle represented non-polar aa changed to polar aa; diamond represented change in charge of polar aa. The substitution hotspots with red fonts represent new identified in this study.

### Spatial distribution characteristic of SARS-CoV-2 in each substitution hotspot

According to the substitution hotspots, we divided two geographical regions, China and outside of China, and studied the distribution differences of mutant and referential type in geographical area at each substitution hotspots, respectively (Fig. 4). As the result showing, except substitution hotspots of ORF1ab-10818 and ORF3a-752, mutant type and referential type in the 12 other substitution hotspots showed significant spatial distribution differences (p<0.01, chi-square test) between China and outside of China. Furthermore, we found in substitution hotspots of ORF1ab-8517 and ORF8-251C, the mutant type (ORF1ab-8517T or ORF8-251C) owned a higher ratio and were more prevalent in China than outside of China, opposite to other 10 substitution hotspots.

**Fig. 4.**
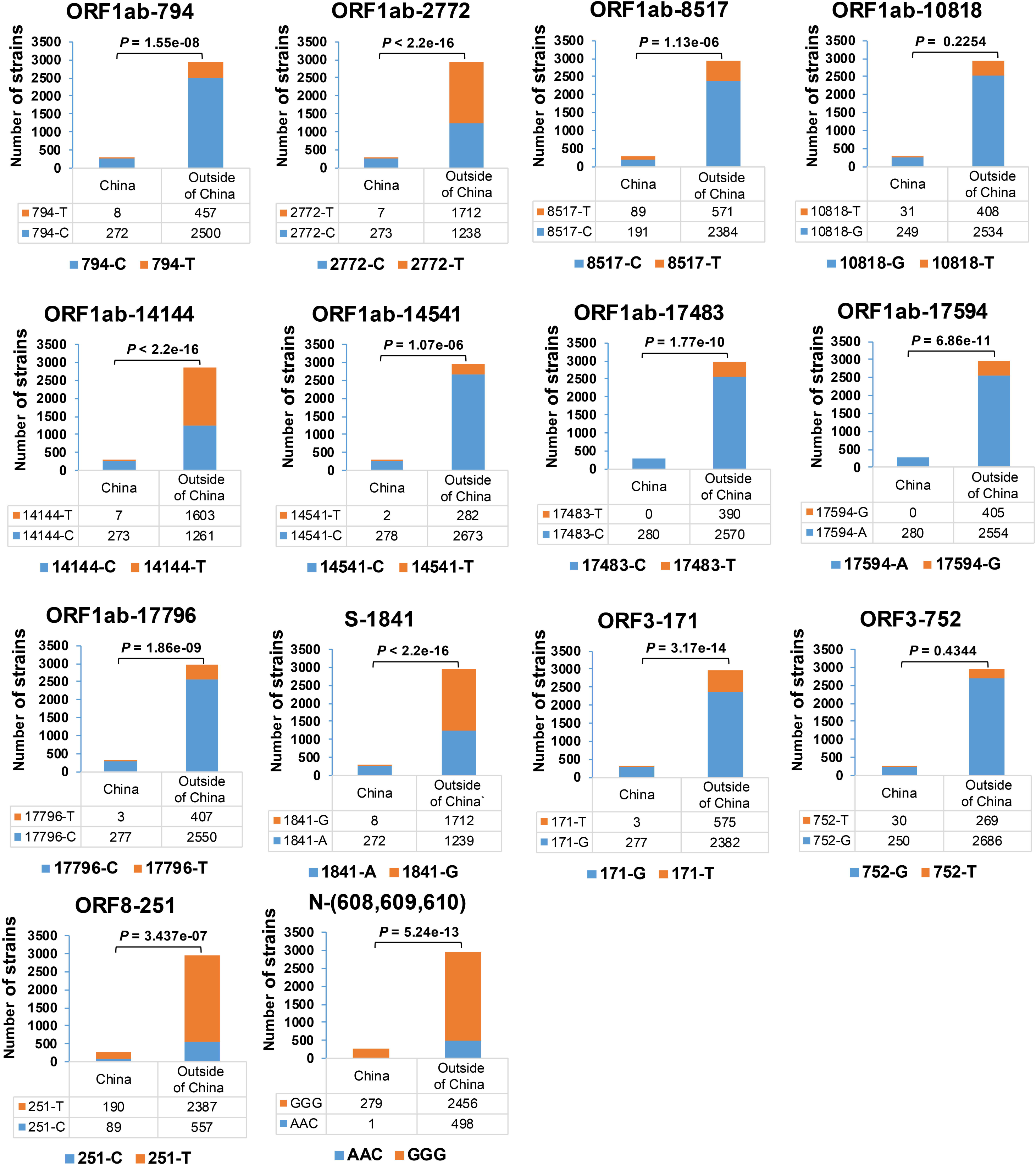
Spatial distribution of referential types and mutant types in 14 substitution hotspots. Note strains which contained degenerate bases in substitution hotspots were culled, and spatial distribution differences of mutant type and referential type in hotspots was based on chi-square test.

### Linkage among these substitution hotspots

Make further efforts, we found some hotspots contains substitutions with similar patterns and frequency (Table 2), indicating potential connection among these substitution hotspots, and then we found 3 groups of possible linkage substitution. In order to facilitate research, we used gene names and base position to mark them, these linkage substitution hotspots were most likely ORF1ab-2772 & ORF1ab-14144 & S-1841, ORF1ab-8517 & ORF8-251 and ORF1ab-17483 & ORF1ab-17594 & ORF1ab-17796.

**Table 2.**
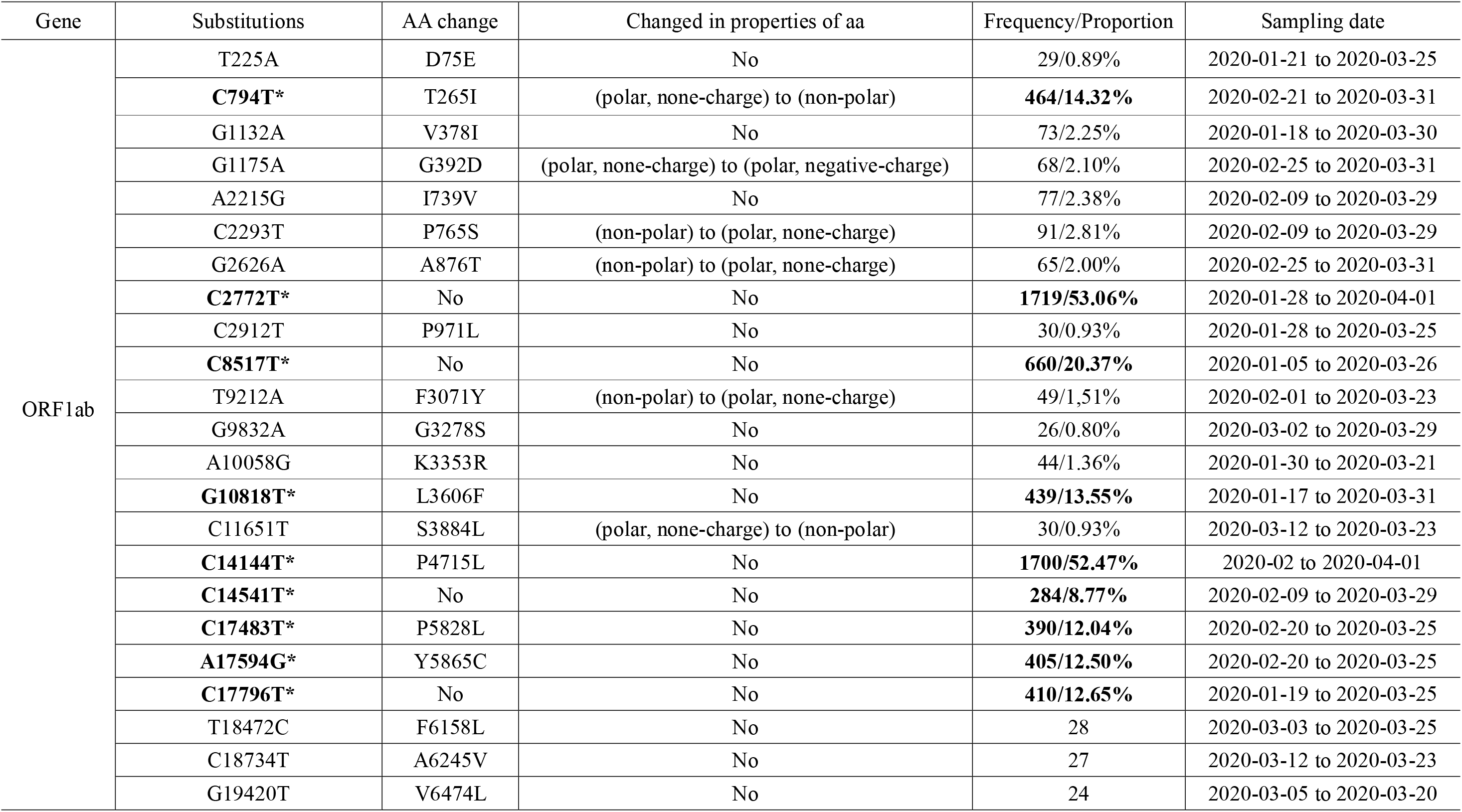
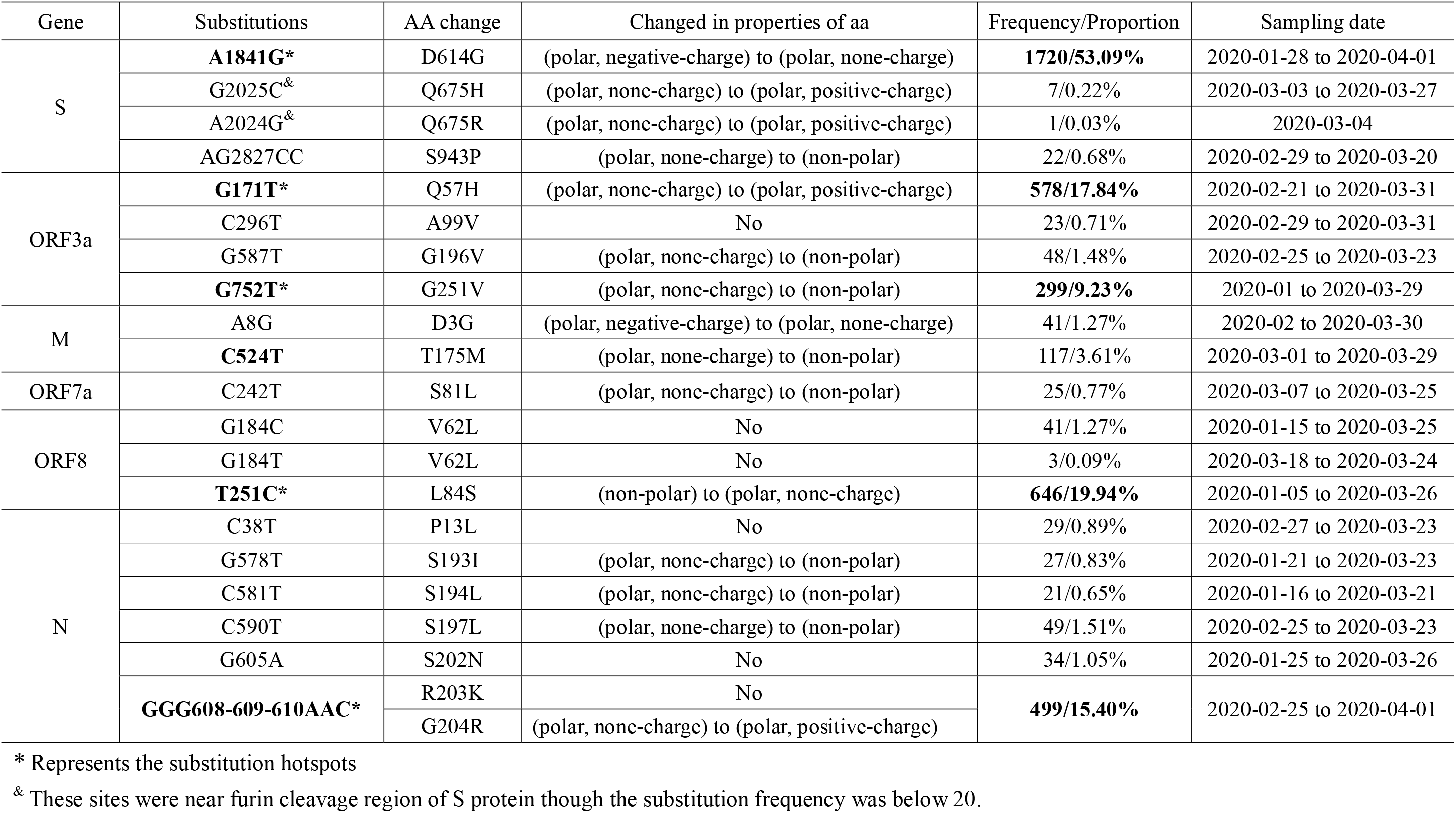
Sites with nonsynonymous substitution frequency over 20 or synonymous substitution frequency over 200

To test our hypothesis, we compared the number of strains between referential type and mutant type which synchronously mutated at the possible linkage substitution hotspots (Table 3). For each combination of possible linkage substitution hotspots, we found mutant type and referential type in accounts for more than 98% of the population, it implied that the genetic variants in these combined substitution hotspots were not independent, but linkage.

**Table 3.**
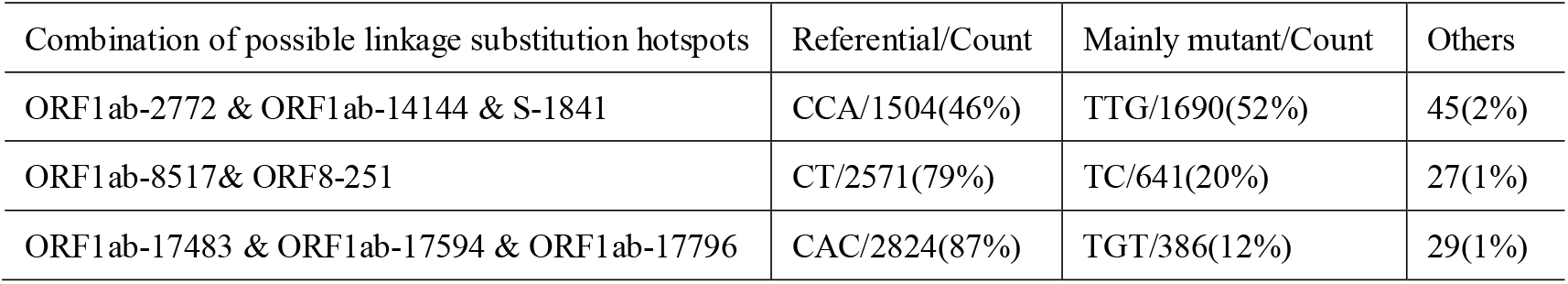
The number of strains with possible linkage substitution among mainly mutant and referential strain

| Combination of possible linkage substitution hotspots | Referential/Count | Mainly mutant/Count | Others |
| --- | --- | --- | --- |
| ORF1ab-2772 & ORF1ab-14144 & S-1841 | CCA/1504(46%) | TTG/1690(52%) | 45(2%) |
| ORF1ab-8517& ORF8-251 | CT/2571(79%) | TC/641(20%) | 27(1%) |
| ORF1ab-17483 & ORF1ab-17594 & ORF1ab-17796 | CAC/2824(87%) | TGT/386(12%) | 29(1%) |

### Vital substitution hotspots in SARS-CoV-2 NSP13

Given 541^th^ aa was one of known key sites of NSP13 for binding to nucleic acids in SARS-CoV, the non-synonymous substitution hotspot of ORF1ab-Y5865C (NSP13-Y541C) in NSP13 probably affects the function of NSP13. We conducted protein model prediction of SARS-CoV-2 NSP13 by homology protein modeling. The result showed that closest protein model to SARS-CoV-2 NSP13 was 6jyt.2.A with 99.83% amino acid identity, which came from SARS-CoV. As the result shown, the tertiary structure of NSP13 were almost completely overlapping between SARS-CoV-2 and SARS-CoV (Fig. 5A). Besides, the 541^th^ aa was relatively conservative in SARS-CoV-2 related animal coronaviruses (Fig. 5B).

**Fig. 5.**
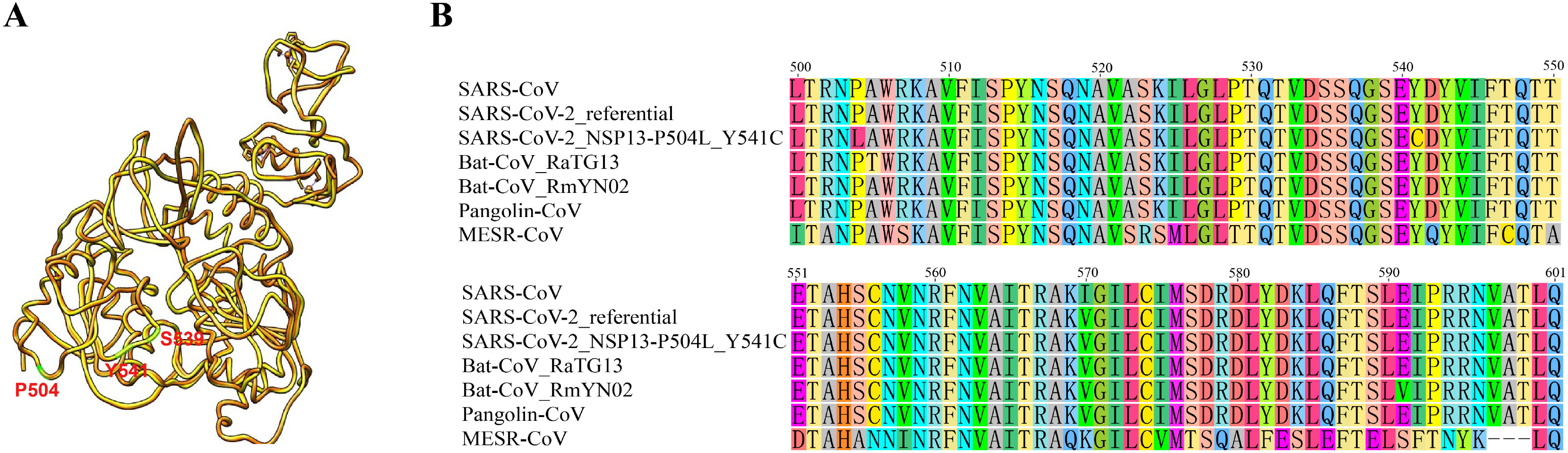
Tertiary structure prediction of SARS-CoV-2 NSP13. (A) Tertiary structure superposition of NSP13 between SARS-CoV-2 (orange) and SARS-CoV (brown, 6jyt.2.A), the tertiary structure of SARS-CoV-2 NSP13 was based on model 6jyt.2.A of SARS-CoV by homology modeling using gene sequence of EPI_ISL_402119. (B) The sequence alignment with partial amino acid of NSP13. SARS-CoV(AY291315.1), SARS-CoV-2_ referential (EPI_ISL_402119), SARS-CoV-2_NSP13-P504L_Y541C (EPI_ISL_413456), Bat-CoV RaTG13(MN996532.1), Bat-CoV_RmYN02 (EPI_ISL_412977), Pangolin-CoV(EPI_ISL_410721), MERS-CoV(KC875821.1).

### Non-synonymous substitutions sites near furin cleavage region

Two non-synonymous substitution sites near the furin cleavage region of PRRA on S protein were identified (Fig. 6, Table 2). There were 7 mutant strains with S-G2025C (S-Q675H) and one with S-A2024G (S-Q675R) among 3239 sequenced strains. 675^th^ aa on S protein was the sixth amino acid upstream of PRRA, and we found S-Q675H and S-Q675R made original polar amino acids from no-charged to positively charged.

**Fig. 6.**
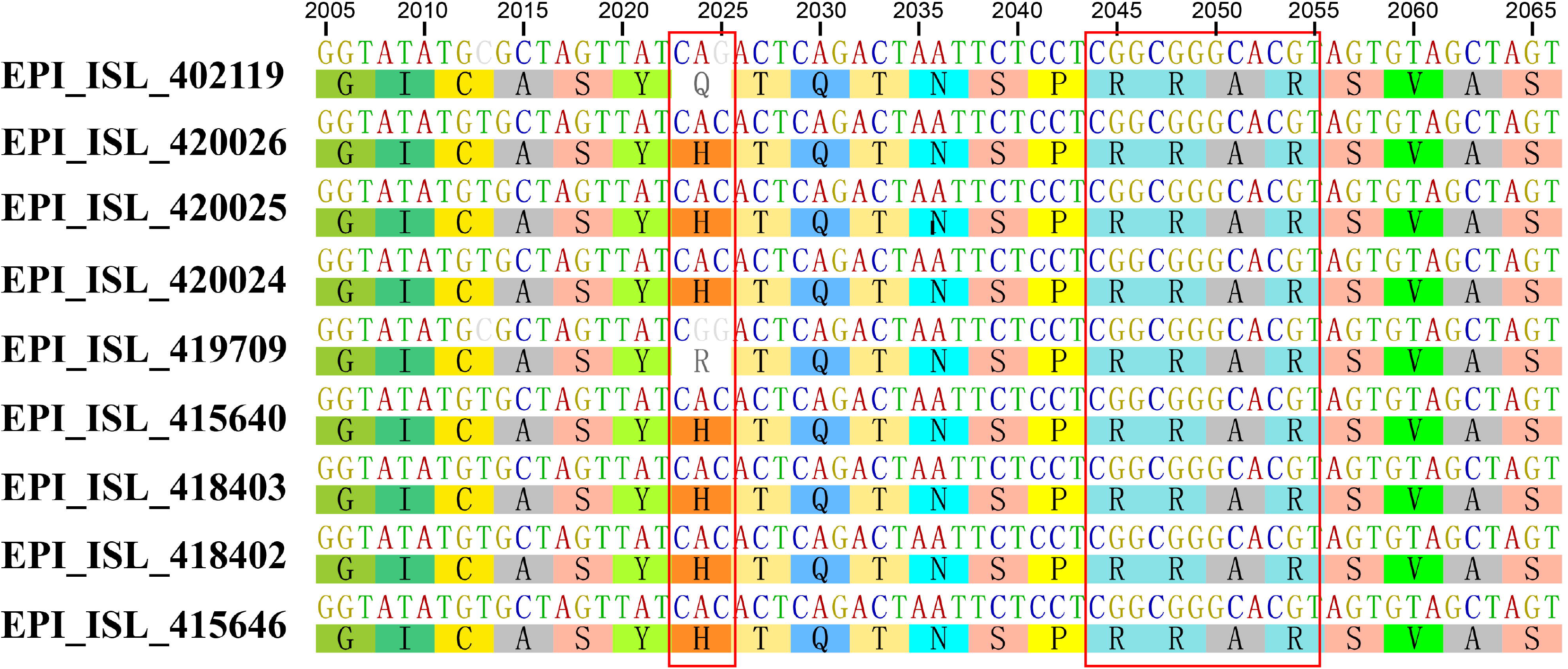
Non-synonymous substitution near the furin cleavage site of S.

### Distinctive polymorphism in SR-rich fragment of SARS-CoV-2

We found a continuous variable area with non-synonymous substitutions from 183th to 204th aa on the N protein in SARS-CoV-2 (Fig. 7), and there were 5 variably sites with non-synonymous substitutions frequency over 20, including a trinucleotide substitution which led to two consecutive aa substitution of R203K and G204R (Table 2). Notably, there were 2 strains substituted on the 196^th^ and 201^th^ codons of SARS-CoV-2, too, but they were synonymous substitutions which did not cause amino acid substitutions (sTable 1).

**Fig. 7.**
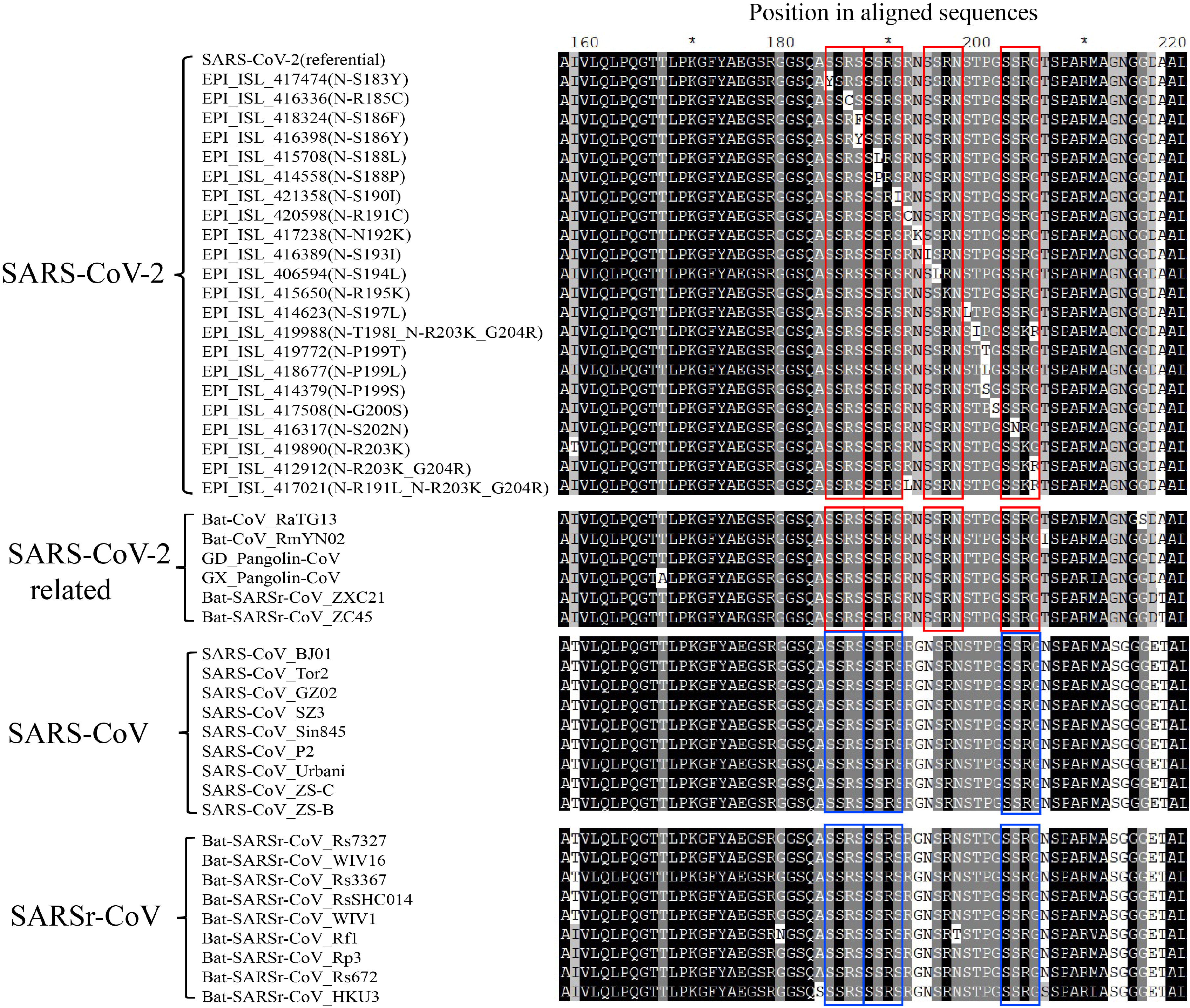
Polymorphic SR-rich region in SARS-CoV-2 N protein. The mutant strain in SARS-CoV-2 was represented using earliest sampled strain of this mutant type. The red and blue boxes represent the SRXX-repeat domain.

The variable area was rich in Ser (S) and Arg (R) and contains SRXX repeat fragments. We compared the region among SARS-CoV-2, SARS-CoV-2 related coronavirus, SARS-CoV and SARSr-CoV in this area (Fig. 7), and found that it was relatively conserved in SARS-CoV-2 related coronavirus, SARSr-CoV and SARS-CoV, but showed distinctive polymorphic in SARS-CoV-2. Interestingly, we found 4 SRXX repeat fragments in most strains of SARS-CoV-2, SARS-CoV-2 related coronavirus strains, but SARSr-CoV and SARS-CoV lacked the third SRXX repeat fragments due to one amino acid substitution. We found the number of SRXX repeat fragments well reflects the evolutionary relationship among SARS-CoV-2, SARS-CoV-2 related coronavirus, SARSr-CoV and SARS-CoV. Especially, he substitution of two continuously amino acids on the last SRXX repeat fragments (203^th^ and 204^th^ in SARS-CoV-2) was exclusively observed in SARS-CoV-2.

To explore the polymorphism of SR-rich region (aa 183-204) on SARS-CoV-2 N protein, we intercepted the amino acids in this region. After culling some strains sequence containing degenerate bases in the region that could not be translated normally, we screened the SR-rich regions in the remaining 3233 strains. The results showed the SR-rich regions in SARS-CoV-2 can be divided into 23 different types (Table 4), including the reference type. The two main types were the reference type (2561 strains) and the mutant type of N-R203K_G204R (494 strains). The majority of the 22 mutant types harbored only single-amino-acid substitution compared to the reference type, indicating that this region was relatively conserved among most SARS-CoV-2 strains. However, we could still find some distinctive polymorphism among different strains. The sampling date showed that most mutations existed in the 3233 recently sequenced strains, implying a constant evolution of the SR-rich region in SARS-CoV-2 genome.

**Table 4.** The 23 types of SR-rich regions in N protein of SARS-CoV-2. Note: The mutation sites are in bold comparing with the reference strain.

| Type | Amino acid sites and mutants |  |  |  |  |  |  |  |  |  |  |  |  |  |  |  |  |  |  |  |  |  |  | First strain | Sampling date |
| --- | --- | --- | --- | --- | --- | --- | --- | --- | --- | --- | --- | --- | --- | --- | --- | --- | --- | --- | --- | --- | --- | --- | --- | --- | --- |
|  | SSXX-1 |  |  |  | SSXX-2 |  |  |  |  |  | SSXX-3 |  |  |  |  |  |  |  | SSXX-4 |  |  |  |  |  |  |
|  | 183 | 184 | 185 | 186 | 187 | 188 | 189 | 190 | 191 | 192 | 193 | 194 | 195 | 196 | 197 | 198 | 199 | 200 | 201 | 202 | 203 | 204 | No. |  |  |
| Reference type | S | S | R | S | S | S | R | S | R | N | S | S | R | N | S | T | P | G | S | S | R | G | 2561 | EPI_ISL_402123 | 2019-12-24 |
| N-R203K_G204R | S | S | R | S | S | S | R | S | R | N | S | S | R | N | S | T | P | G | S | S | K | R | 494 | EPI_ISL_412912 | 2020-02-25 |
| N-S197L | S | S | R | S | S | S | R | S | R | N | S | S | R | N | L | T | P | G | S | S | R | G | 49 | EPI_ISL_414623 | 2020-02-25 |
| N-S202N | S | S | R | S | S | S | R | S | R | N | S | S | R | N | S | T | P | G | S | N | R | G | 34 | EPI_ISL_416317 | 2020-01-25 |
| N-S193I | S | S | R | S | S | S | R | S | R | N | I | S | R | N | S | T | P | G | S | S | R | G | 27 | EPI_ISL_416389 | 2020-01-21 |
| N-S194L | S | S | R | S | S | S | R | S | R | N | S | L | R | N | S | T | P | G | S | S | R | G | 21 | EPI_ISL_406594 | 2020-01-16 |
| N-S188L | S | S | R | S | S | L | R | S | R | N | S | S | R | N | S | T | P | G | S | S | R | G | 6 | EPI_ISL_415708 | 2020-03-07 |
| N-S183Y | Y | S | R | S | S | S | R | S | R | N | S | S | R | N | S | T | P | G | S | S | R | G | 6 | EPI_ISL_417474 | 2020-03-12 |
| N-R195K | S | S | R | S | S | S | R | S | R | N | S | S | K | N | S | T | P | G | S | S | R | G | 6 | EPI_ISL_415650 | 2020-03-02 |
| N-R185C | S | S | C | S | S | S | R | S | R | N | S | S | R | N | S | T | P | G | S | S | R | G | 5 | EPI_ISL_416336 | 2020-02-01 |
| N-S190I | S | S | R | S | S | S | R | I | R | N | S | S | R | N | S | T | P | G | S | S | R | G | 4 | EPI_ISL_421358 | 2020-03-17 |
| N-S188P | S | S | R | S | S | P | R | S | R | N | S | S | R | N | S | T | P | G | S | S | R | G | 4 | EPI_ISL_414558 | 2020-03-06 |
| N-R191L & N-R203K_G204R | S | S | R | S | S | S | R | S | L | N | S | S | R | N | S | T | P | G | S | S | K | R | 4 | EPI_ISL_417021 | 2020-03-15 |
| N-G200S | S | S | R | S | S | S | R | S | R | N | S | S | R | N | S | T | P | S | S | S | R | G | 2 | EPI_ISL_417508 | 2020-03-16 |
| N-N192K | S | S | R | S | S | S | R | S | R | K | S | S | R | N | S | T | P | G | S | S | R | G | 2 | EPI_ISL_417238 | 2020-03-03 |
| N-P199S | S | S | R | S | S | S | R | S | R | N | S | S | R | N | S | T | S | G | S | S | R | G | 1 | EPI_ISL_414379 | 2020-02-18 |
| N-P199T | S | S | R | S | S | S | R | S | R | N | S | S | R | N | S | T | T | G | S | S | R | G | 1 | EPI_ISL_419772 | 2020-03-13 |
| N-P199L | S | S | R | S | S | S | R | S | R | N | S | S | R | N | S | T | L | G | S | S | R | G | 1 | EPI_ISL_418677 | 2020-03-15 |
| N-T198I & N-R203K_G204R | S | S | R | S | S | S | R | S | R | N | S | S | R | N | S | I | P | G | S | S | K | R | 1 | EPI_ISL_419988 | 2020-03-23 |
| N-R203K | S | S | R | S | S | S | R | S | R | N | S | S | R | N | S | T | P | G | S | S | K | G | 1 | EPI_ISL_419890 | 2020-03-19 |
| N-S186F | S | S | R | F | S | S | R | S | R | N | S | S | R | N | S | T | P | G | S | S | R | G | 1 | EPI_ISL_418324 | 2020-03-11 |
| N-R191C | S | S | R | S | S | S | R | S | C | N | S | S | R | N | S | T | P | G | S | S | R | G | 1 | EPI_ISL_420598 | 2020-03-22 |
| N-S186Y | S | S | R | Y | S | S | R | S | R | N | S | S | R | N | S | T | P | G | S | S | R | G | 1 | EPI_ISL_416398 | 2020-02-02 |

## Discussion

Program tools for high-throughput analyses on big sequencing data of genome are of great importance for genomic and evolutionary studies. In comparative genomic studies, it is very important to identify whether amino acid properties have changed when dealing with non-synonymous substitutions, which can help to identify some important mutations. More importantly, simplicity of operation and high summary of analysis results could save a lot of time for researchers. Thus, we have developed the BioAider program, aiming to liberate researchers from numerous sequencing data. BioAider V1.0 contains more than 10 functional programs for sequence processing and analysis. It can be applied to assisting gene batch annotation and quickly mutations analysis for coding gene. For mutations analysis, BioAider can distinguish and summarize all the nucleotide variations sites with corresponding mutation type and frequency, and then generates the frequency distribution map of synonymous and non-synonymous substitution sites. The expansion and optimization of BioAider in the future will endow the software more functions.

In this study, we conducted high-resolution analysis of mutations within SARS-CoV-2 genome based on 3240 sequenced strains using BioAider. 2152 mutation sites among different strains were detected which accounted for 7.36% of the complete genome. These mutation sites included 1,335 non-synonymous substitution sites, implying abundant variations in the genome of SARS-CoV-2. However, more than half of the mutation sites were only observed in a single strain of SARS-CoV-2 which was difficult to be distinguished from sequencing errors. Considering there were only 60 sites with substitution frequency above 20. Therefore, we speculated that there were no large-scale mutations in the sequencing strains we analyzed.

According to the spatial distribution analysis between strains of mutant types and referential type regarding the substitution hotspots, 12 of 14 sites showed significant distribution differences in China and outside of China, indicating different virus prevalence in the two regions. We also noticed mostly strains sampled in China were before Mar 2020, this was due to the epidemic of China has been roughly controlled in March. Considering these facts, we speculate that virus differentiation in and outside of China may attribute to human intervention.

In SARS-CoV NSP13, the 541^th^ aa is critical for the protein function and double mutations of S539A/Y541A show higher unwinding activity for nucleic acids binding [17]. The amino acid identity of NSP13 between SARS-CoV-2 and SARS-CoV was 99.83% and their tertiary structure was almost completely overlapping, indicating that NSP13 541^th^ aa is also a vital site for the function of SARS-CoV-2 NSP13. Notably, the mutant strain of NSP13-541C first appeared in the United States (sampled on Feb 20, 2020), and the United States was the country with the largest number and extremely high proportion of this mutant strains (316, 78% of all the 405 mutant strains with NSP13-Y541C). We also noted that there were a total of 711 strains sampled in the United States on Feb 20, 2020 and beyond, strains with NSP13-Y541C accounted for almost half of these strains in the United States. This fact indicates that NSP13-Y541C has gone through less negative selection pressure and even this variation may be beneficial. Therefore, we speculate that NSP13-Y541C improves the unwinding activity of NSP13 and promotes the replication of SARS-CoV-2, contributing to their rapid spread in the United States. However, further studies are required to reveal the detailed effect of substitution in 541^th^ aa of SARS-CoV-2 NSP13 and whether the possible linkage substitution site of 504^th^ aa plays a synergistic role in this process.

Among the potential linkage substitution hotspots identified in this study, ORF1ab-8517 and ORF8-251 were observed in a recent research [31, 34]. Among the triple linkage substitution hotspots of ORF1ab-277, ORF1ab-14144 and S-1841, ORF1ab-C2772T belongs to synonymous substitution, while ORF1ab-C14144T caused the aa change of NSP12-P323L in interface region which is a bridge section connecting NiRNA (nidovirus RdRp-associated nucleo-tidyltransferase) and Fingers of RdRp [14]. The 614^th^ aa in S protein was located in the subdomain (SD) region downstream of the receptor-binding domain (RBD) on S1. We found that mutant strains with NSP12-323L & S-614G were popular in the world and owned more than half the frequency (52%) in the population, indicating that this mutant was dominant in SARS-CoV-2. However, the functional impact of amino acid changed in these two sites is still unclear at present, and whether NSP12-323 and S-614 are related on specific function needed more experimental data to verify.

Previous studies have reported that SR-rich region in SARS-CoV is crucial for N protein multimerization and the interaction with membrane (M) protein [35, 36]. In the SR-rich region of N protein, there was only 2-aa difference between SARS-CoV-2 (referential type) and SARS-CoV. Therefore, the similar function of SR-rich region may exist in SARS-CoV-2, although there is no relevant research reported at present. Compared to SARS-CoV, SARS-CoV-2 harbors one more SRXX repeat fragments in the 193^th^ −196^th^ aa of N protein. Previous research reported that the SR-rich region of SARS-CoV in 184^th^ −196^th^ aa was crucial for N protein multimerization and the deletion of this region completely would make N protein abolish the self-multimerization [36]. Besides, SRXX repeat fragments may play an important role in SARS-CoV infection [37]. We noted that SARS-CoV could be regarded as a deletion of SRXX in SR-rich region compared to SARS-CoV-2, and the transmission rate of SARS-CoV-2 was higher than SARS-CoV, thus, whether SR-rich region of SARS-CoV-2 plays an important role in this process is worth to explore. However, the research about SR-rich region in coronavirus was still limited at present. Especially, unlike SARS-CoV, the region in SARS-CoV-2 is still constantly evolving, implying that the N protein of SARS-CoV-2 may employ a more flexible replication mechanism and even interaction with M protein. A study reported that the N protein of SARS-CoV can specifically bind to heterogeneous nuclear ribonucleoproteins (hnRNPs) A1 and plays an important role in RNA replication and transcription, especially, the key binding region is in the SR-rich region of SARS-CoV N protein (aa 161-210), the interaction between human hnRNP A1 and SARS-N protein may be the key to SARS-CoV replication and transcription [37]. Whether such a similar mechanism exists in SARS-CoV-2 is still unclear. However, in human cells, there are more than 20 hnRNPs has been discovered, and SR-rich region of SARS-CoV-2 showing distinctive polymorphism, whether the SARS-CoV-2 can bind to one of these hnRNPs needs more verification. It might provide a new hint in understanding the process of SARS-CoV-2 replication in human cells. Besides, the potential phenotype changes related to nonsynonymous substitution hotspots on third SRXX repeat fragments of SARS-CoV-2 may be also worth attention.

In the hotspot screening, we have found several substitutions critical for viral replication potentially. For instance, the ORF1ab-C794T was located on NSP2 and NSP2 was reported to inhibit the host protein PHB1 and PHB2 which benefited viral replication, indicating that ORF1ab-C794T may affect viral replication [11]. The ORF3a-G171T (Q57H) was located at the transmembranous domain of the 3a protein (sFig. 2). 3a protein was reported to form ion channels on host cell membrane and enhance the membrane permeability which benefited SARS-CoV life cycle [38]. Thus, ORF3a-G171T (Q57H) may affect the formation of ion channels and subsequently influence the viral replication. As for ORF3a-G752T (G251V), a recent analysis predicted that it was in an important functional domain and might be related to virulence, infectivity, ion channel formation and virus release [30]. We also found several important sites for virus entry, though they showed lower substitution frequency in our research data. For instance, S-Q675H and S-Q675R near furin cleavage region possibly influence the cleavage of RRAR, a critical step for virus entry (Fig. 6). We also detected a strain with the mutation of S-R408I located in the RBD (sTable 1) which was reported to play an important role in virus-receptor binding by a recent study [39].

In summary, we initially revealed the variation characteristics of SARS-CoV-2 and predicted the possible impact for some non-synonymous substitution hotspots. The NSP13-Y541C was crucial, which may affect the unwinding activity of NSP13 and the replication of SARS-CoV-2. Especially, the SR-rich region of N protein in SARS-CoV-2 shows distinctive polymorphism, and the quantity of SRXX repeat fragments well reflects the evolutionary relationship among SARS-CoV-2, SARS-CoV-2 related coronavirus, SARSr-CoV and SARS-CoV, it providing a new clue to study the formation mechanism of SARS-CoV-2. In this study, we could not include all the sequences of SARS-CoV-2 due to the rapid increase of the genomic data. More researches or real time analysis with relevant clinical data may contribute to the viral epidemiology of SARS-CoV-2 and treatment of COVID-19.

## Methods and materials

### Main functions and working principle of BioAider

BioAider V1.0 was developed based on Python 3.7 and R 3.5.2, and used PyQt5 for interface packaging. BioAider is a local software containing more 10 subroutines and can run on Windows systems at present (https://github.com/ZhijianZhou01/BioAider/releases).

BioAider is mainly used to process and analysis genome sequencing data (such as viral genome data), the main two functions are mutation analysis and batch extract gene for auxiliary sequence annotation at present. First, for batch processing genes, users can import the aligned complete genome sequence set (*.fasta or *.fas format), and adjust the reference sequence for gene extraction to the forefront of the sequence set. Paste the related gene information of reference sequence in the input box, gene name, star string and end string, which separated by “,” (sFig. 3A). Then user can batch extract genes from a large number of sequences. Note that, the length of start string or end string is not limited in, but it is required to be unique in the reference sequence.

Second, for mutation analysis, BioAider will scan all the codons in the aligned sequence, using the first sequence in the data set as a reference sequence. BioAider can identify five different mutation types (synonymous, non-synonymous, insert, deletion and termination) based on the standard codon method. It can distinguish the changes in properties of amino acid when dealing non-synonymous substitutions. Then, it will locate the position of the corresponding base in the mutated codon, so it can identify multiple types of mutation at the same site, and also well discriminate dinucleotide and trinucleotide substitution. Next, it summarizes all the mutation sites with corresponding frequency and strains, and if user chooses to generate the frequency distribution of synonymous or non-synonymous sites, BioAider will directly generate the results in vector format image. (sFig. 3B). Of note, the frequency distribution map does not include those sites that are both synonymous and non-synonymous substitution, because such sites cannot determine the common substitution frequency.

### Mutation analysis of SARS-CoV-2

The 3240 complete genome sequences of SARS-CoV-2 with relatively higher quality of sequencing were downloaded from GISAID (https://www.gisaid.org/), and the reference genome sequence of SARS-CoV-2 (NC_045512.2) for ORF annotation was from GenBank (https://www.ncbi.nlm.nih.gov/genbank). All the viral strains used in this study were listed in sTable3. Multiple sequence alignment of genomic sequences of SARS-CoV-2 were accomplished using MAFFT v7.407 [40]. The annotation and extraction of codon genes of these 3240 SARS-CoV-2 genome sequences using BioAider V1.0. We extracted 11 continuously coding genes based on the annotation information of NC_045512.2, including ORF1ab, S, ORF3a, E, M, ORF6, ORF7a, ORF7b, ORF8, N and ORF10. Then we used MUSCLE program in MEGA v7.0.14 to align these coding genes based on codons method [41].

We combined these 11 continuously coding genes to tandem sequence in BioAider and used it to represent the complete genome sequence of SARS-CoV-2 for subsequent analysis. Among all the early sampled virus strains (Dec 24, 2019 to Dec30, 2019), genome of Wuhan/IVDC-HB-01/2019 (GISAID ID: EPI_ISL_402119, sampled in Dec30, 2019) occupied most frequently (65 strains sequence were same with IVDC-HB-01/2019) in 3240 sequenced strains and appears in multiple regions, therefore, the strain of EPI_ISL_402119 was used the reference sequence for genome variation analysis of SARS-CoV-2 in BioAider V1.0.

### Structure prediction of protein

The tertiary protein structure was downloaded from the Protein Data Bank (PDB, http://www.rcsb.org), and protein homology modeling was calculated by online tools SWISS-MODEL then using Chimera 1.10.2 to simulate the effect of a single amino acid mutation on protein structure[42, 43].

## Supporting information

Supplemental material summary

