## Supplemental Figure 1 for "Characterization of the substitution hotspots in SARS-CoV-2 genome using BioAider and detection of a SR-rich region in N protein providing further evidence of its animal origin"

ORF1ab

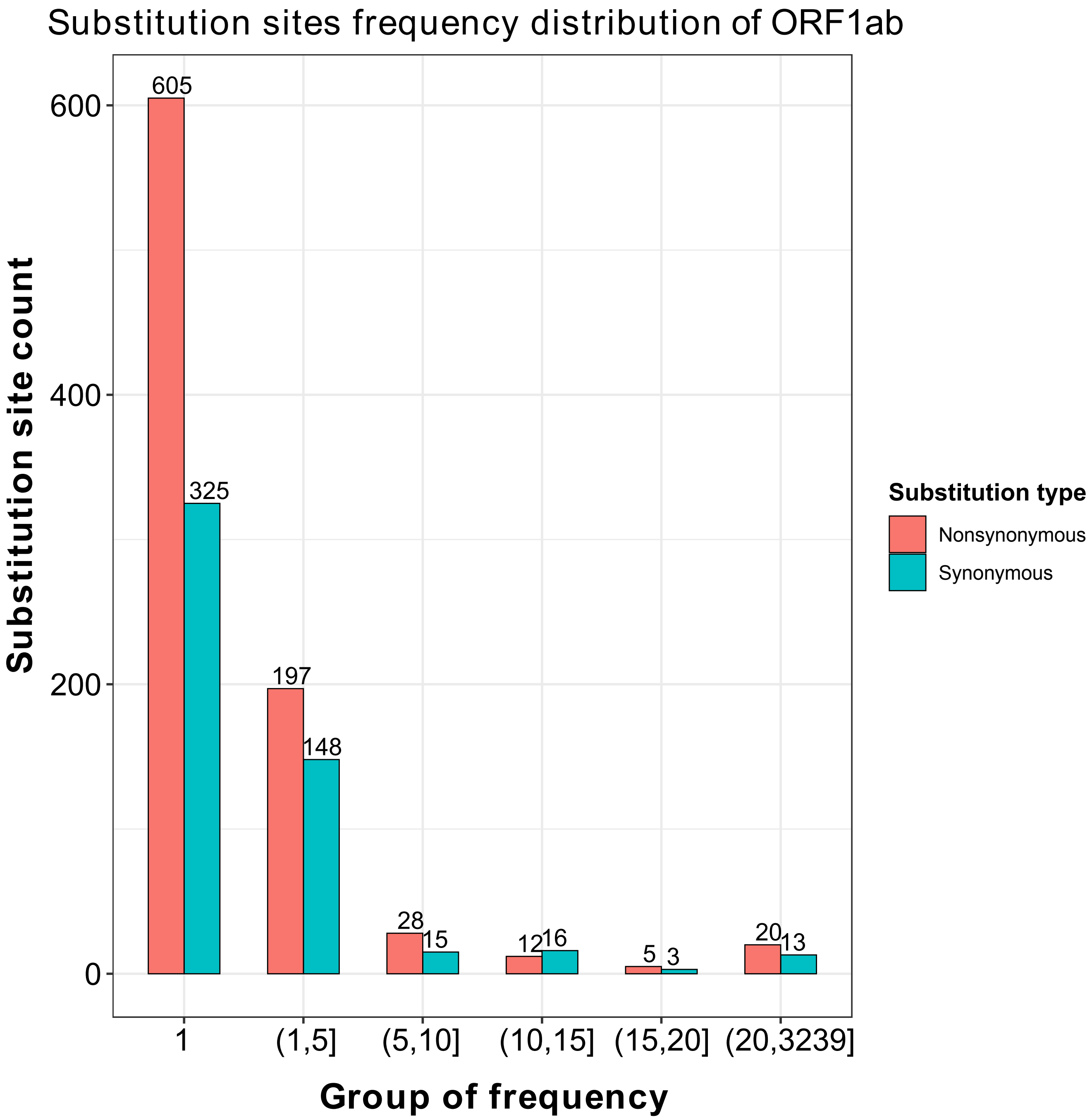

Substitution sites frequency distribution of ORF1ab
Substitution site count
600
400
200
0
605
325
197
148
28
15
12
16
5
3
20
13
1
(1,5]
(5,10]
(10,15]
(15,20]
(20,3239]
Group of frequency
Substitution type
Nonsynonymous
Synonymous

S

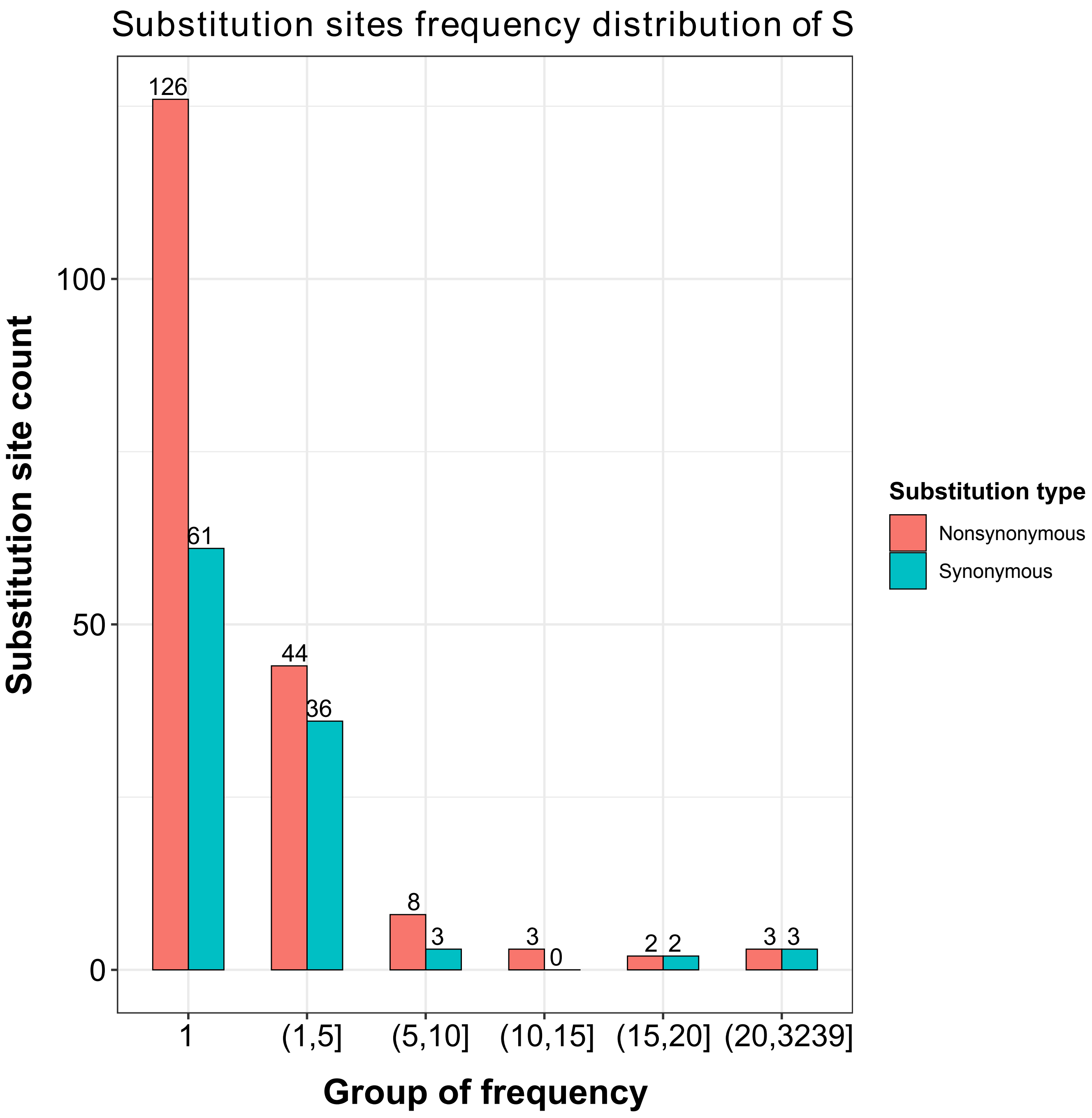

Substitution sites frequency distribution of S
Substitution site count
Group of frequency
Substitution type
Nonsynonymous
Synonymous
0
50
100
1
(1,5]
(5,10]
(10,15]
(15,20]
(20,3239]
126
61
44
36
8
3
3
0
2
2
3
3

ORF3a

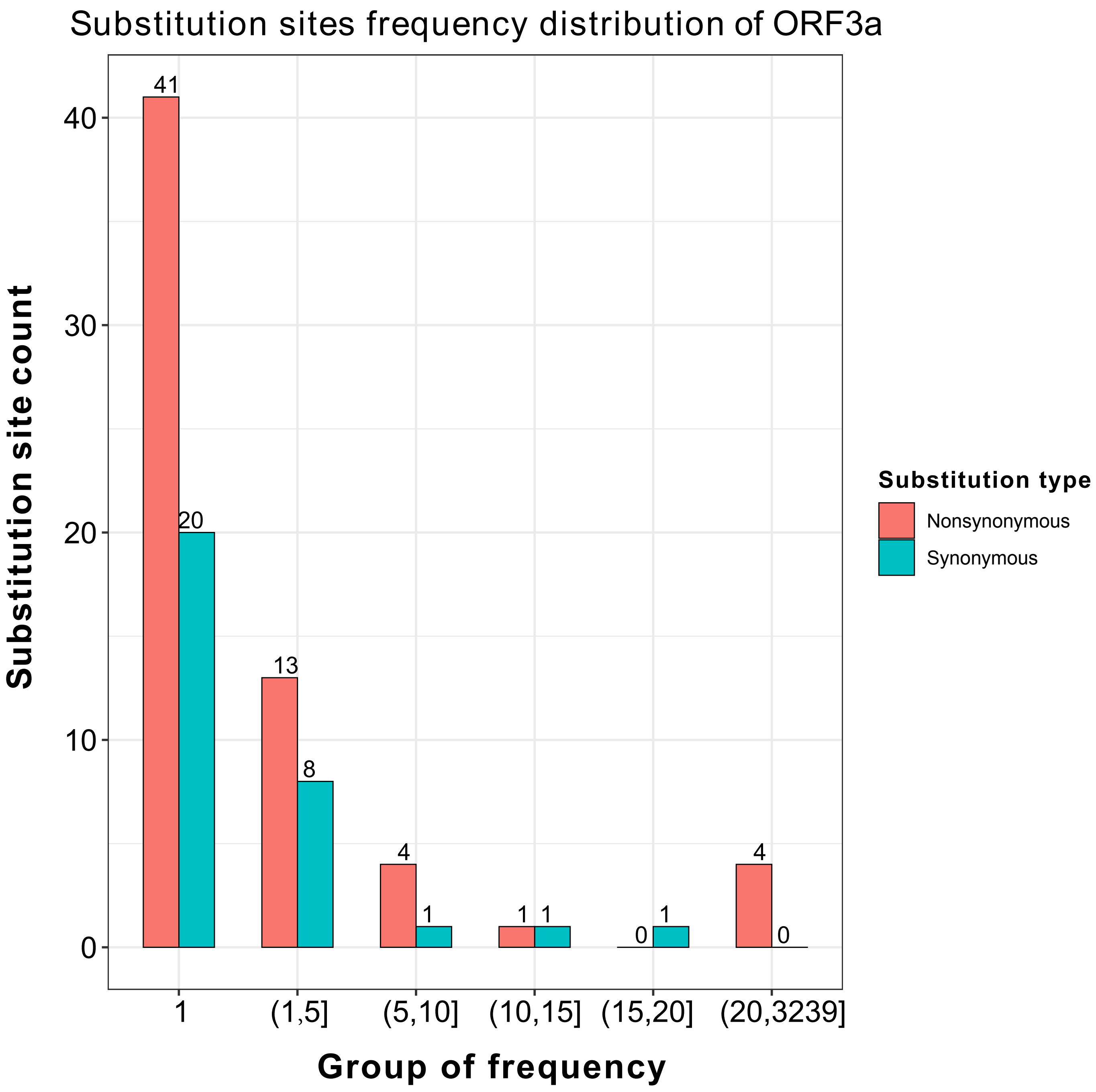

Substitution sites frequency distribution of ORF3a
Substitution site count
Group of frequency
Substitution type
Nonsynonymous
Synonymous
41
20
13
8
4
1
1
1
0
1
4
0
40
30
20
10
0
1
(1,5]
(5,10]
(10,15]
(15,20]
(20,3239]

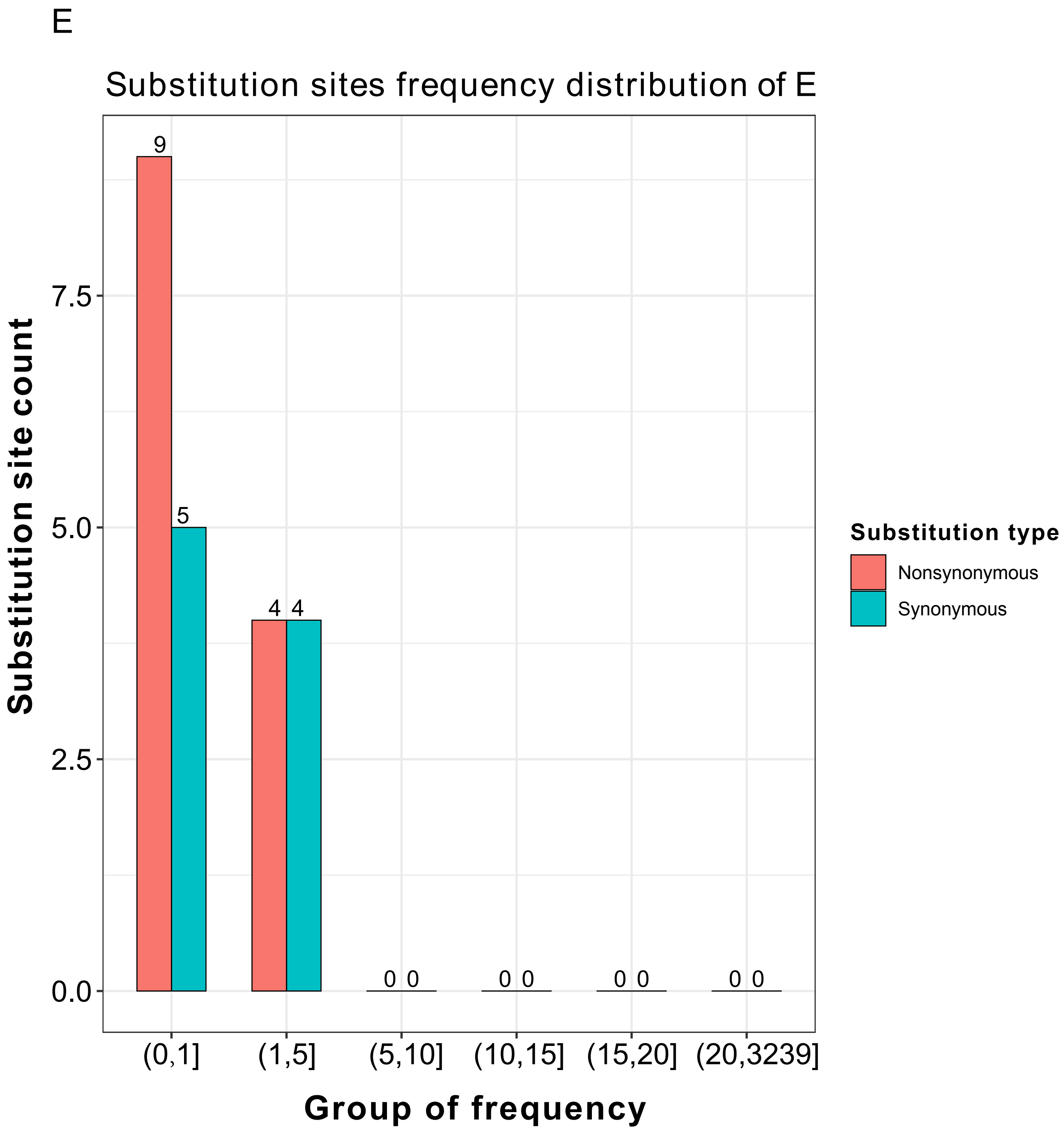
E
Substitution sites frequency distribution of E
Substitution site count
Group of frequency
0.0
2.5
5.0
7.5
9
5
4 4
0 0
0 0
0 0
0 0
(0,1]
(1,5]
(5,10]
(10,15]
(15,20]
(20,3239]
Substitution type
Nonsynonymous
Synonymous

M

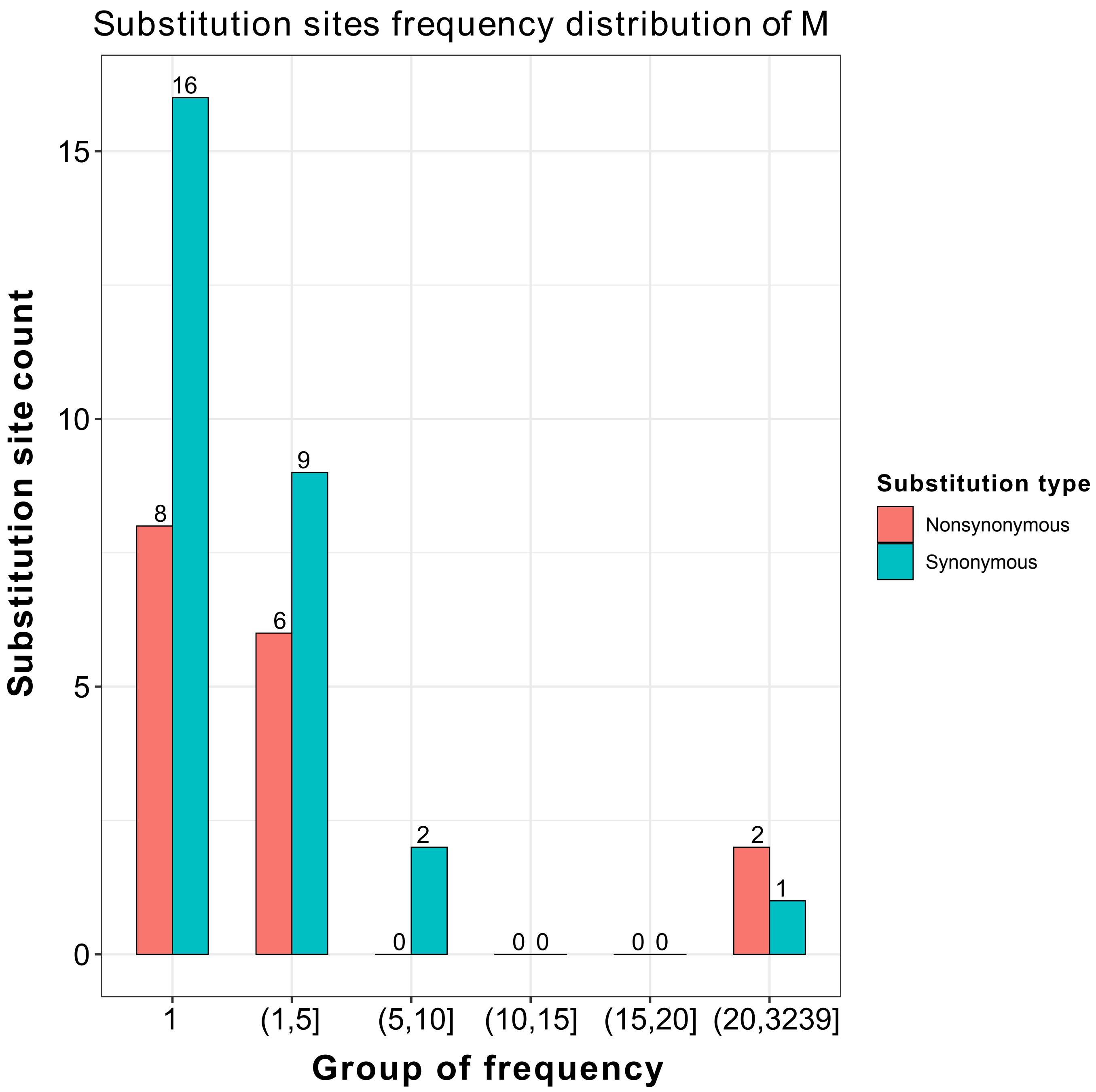

Substitution sites frequency distribution of M
Substitution site count
Group of frequency
Substitution type
Nonsynonymous
Synonymous
0
5
10
15
1
(1,5]
(5,10]
(10,15]
(15,20]
(20,3239]
16
8
9
6
2
0
0 0
0 0
2
1

ORF6

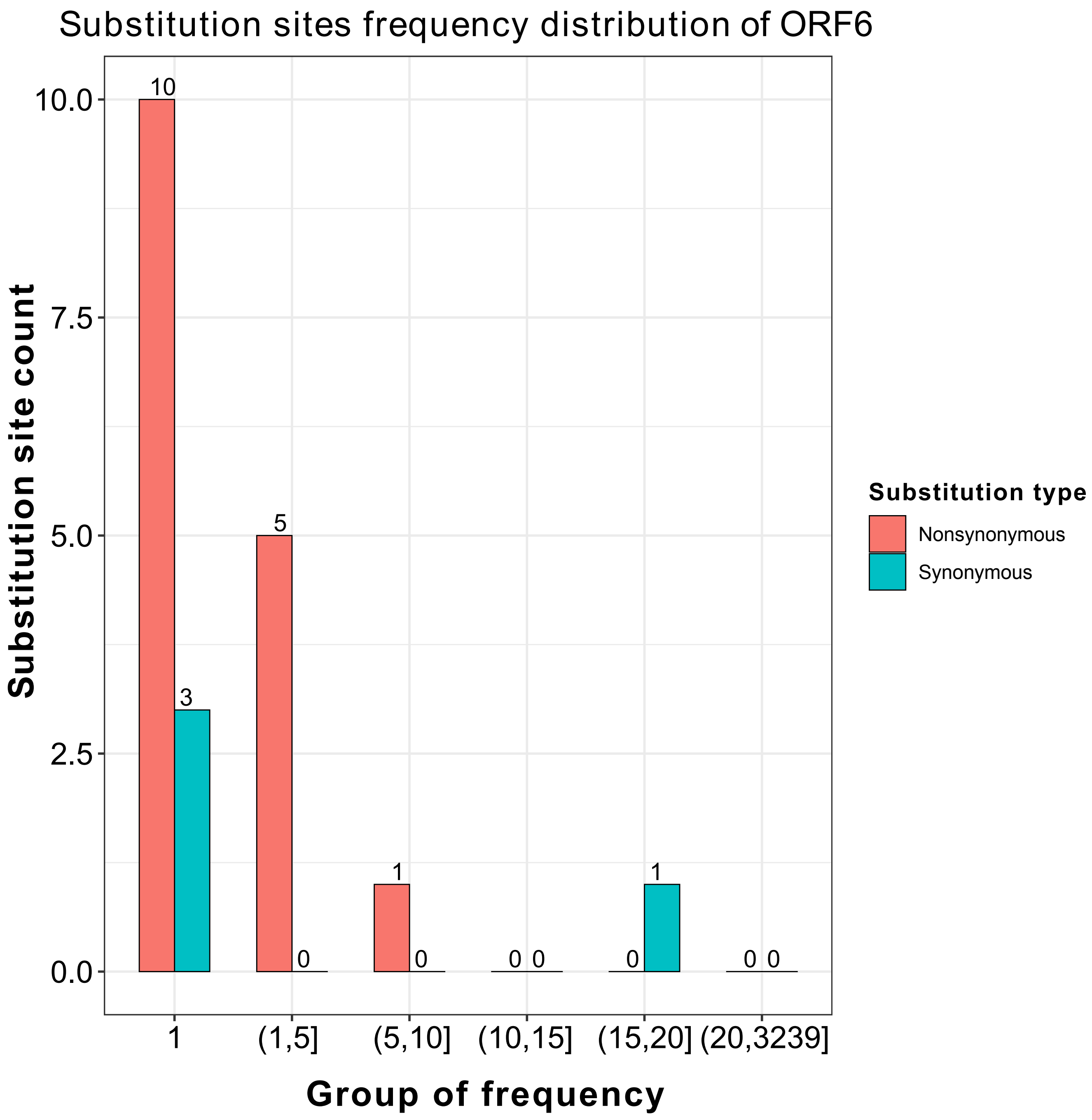

Substitution sites frequency distribution of ORF6
Substitution site count
10.0
7.5
5.0
2.5
0.0
10
3
5
0
1
0
0 0
0
1
0 0
1
(1,5]
(5,10]
(10,15]
(15,20]
(20,3239]
Group of frequency
Substitution type
Nonsynonymous
Synonymous

ORF7a

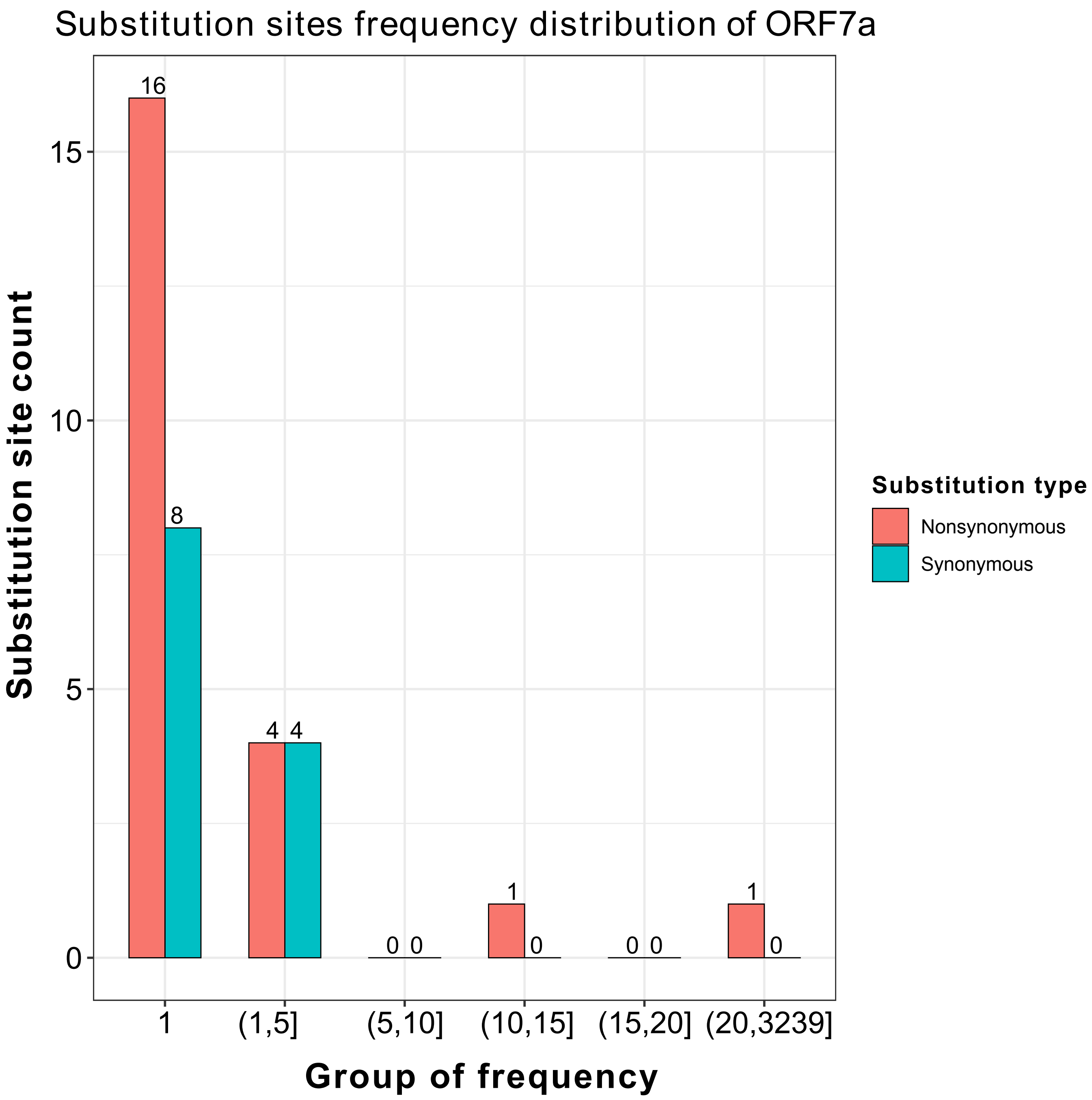

Substitution sites frequency distribution of ORF7a
Substitution site count
Group of frequency
16
8
4
4
0
0
1
0
0
0
1
0
0
5
10
15
1
(1,5]
(5,10]
(10,15]
(15,20]
(20,3239]
Substitution type
Nonsynonymous
Synonymous

ORF7b

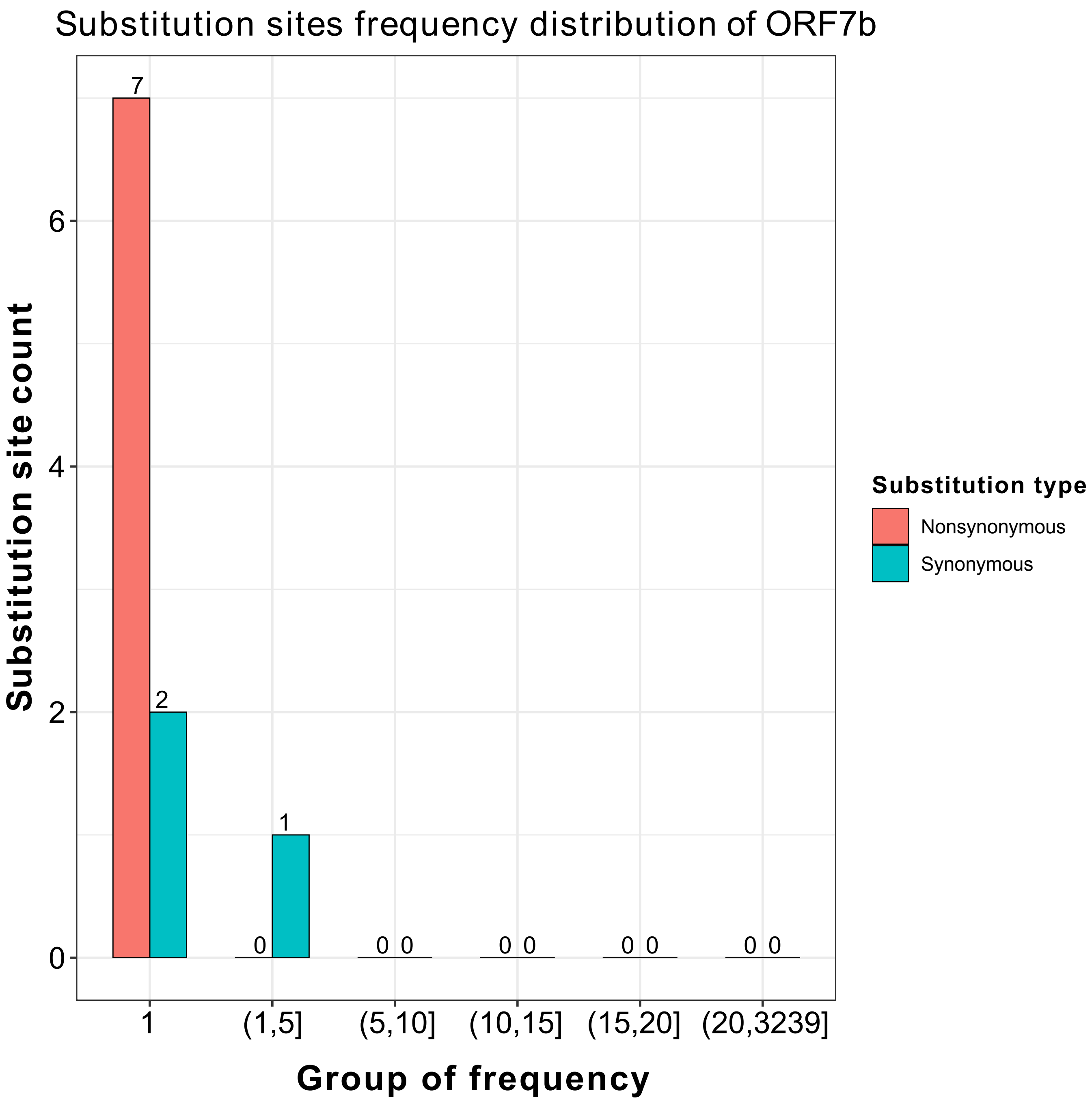

Substitution sites frequency distribution of ORF7b
Substitution site count
Group of frequency
1
(1,5]
(5,10]
(10,15]
(15,20]
(20,3239]
0
2
4
6
7
2
0
1
0 0
0 0
0 0
0 0
Substitution type
Nonsynonymous
Synonymous

ORF8

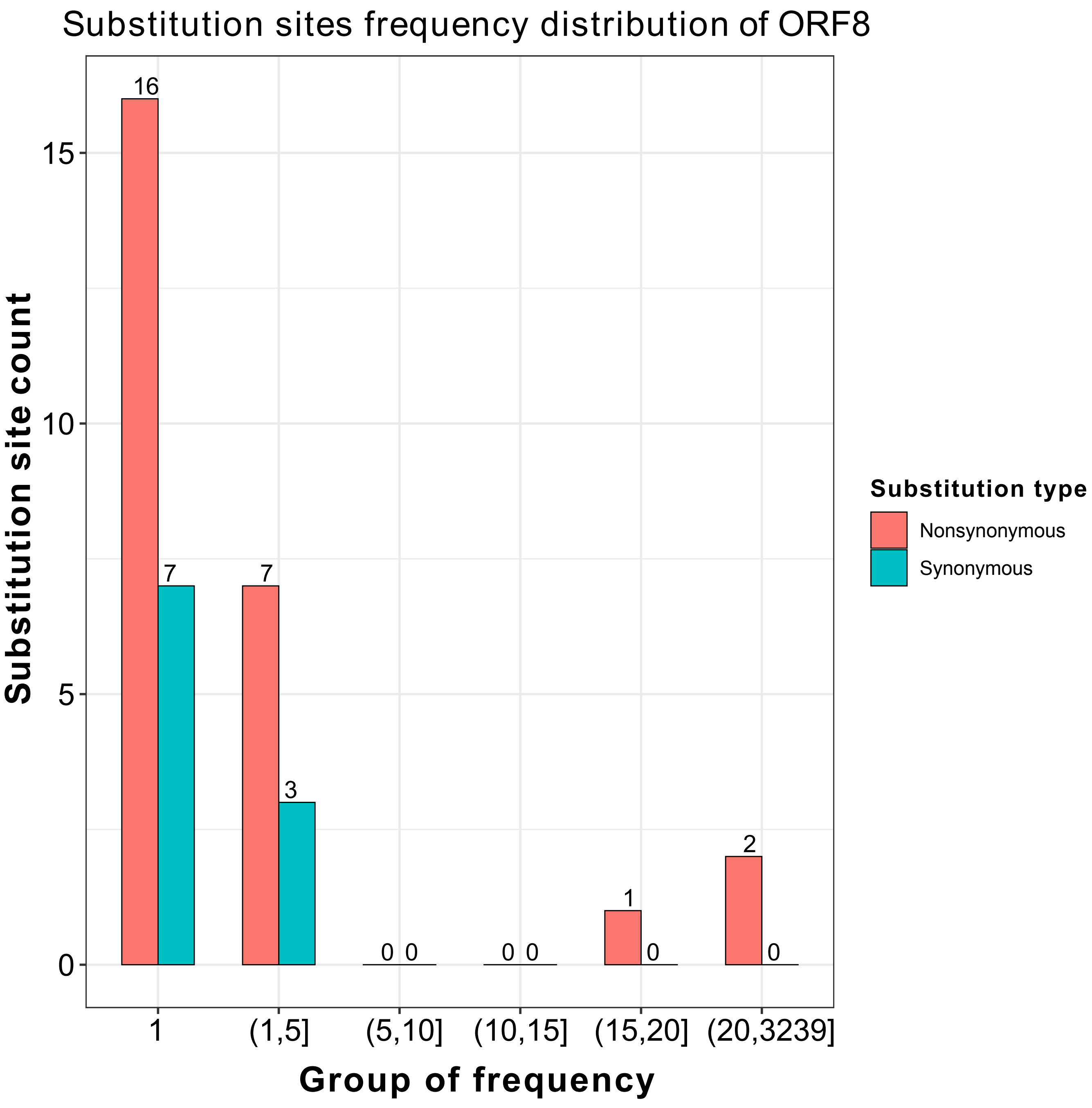
Substitution sites frequency distribution of ORF8
Substitution site count
Group of frequency
1
(1,5]
(5,10]
(10,15]
(15,20]
(20,3239]
0
5
10
15
16
7
7
3
0 0
0 0
1
0
2
0
Substitution type
Nonsynonymous
Synonymous

N

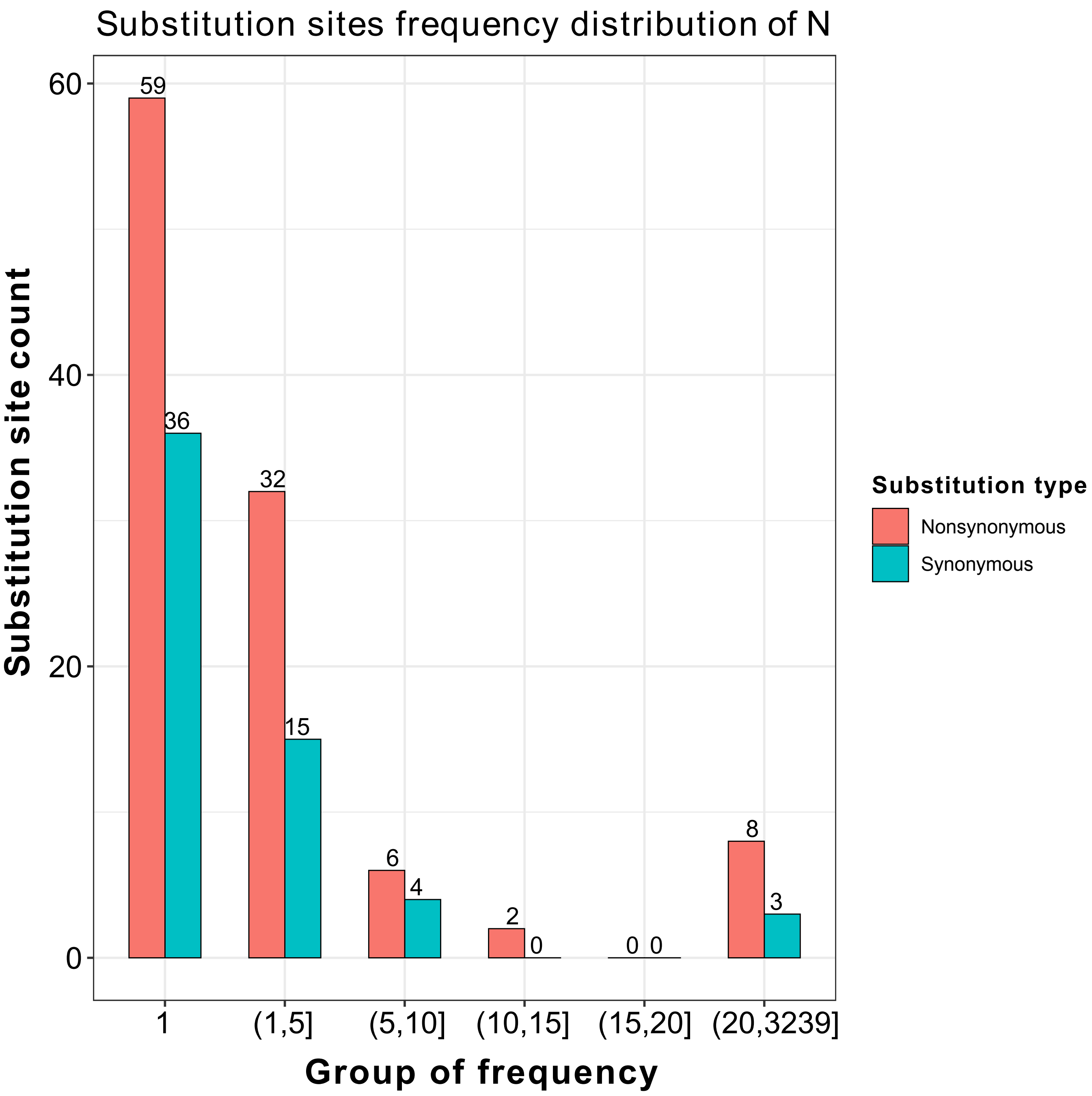

Substitution sites frequency distribution of N
Substitution site count
Group of frequency
60
40
20
0
59
36
32
15
6
4
2
0
0 0
8
3
1
(1,5]
(5,10]
(10,15]
(15,20]
(20,3239]
Substitution type
Nonsynonymous
Synonymous

ORF10

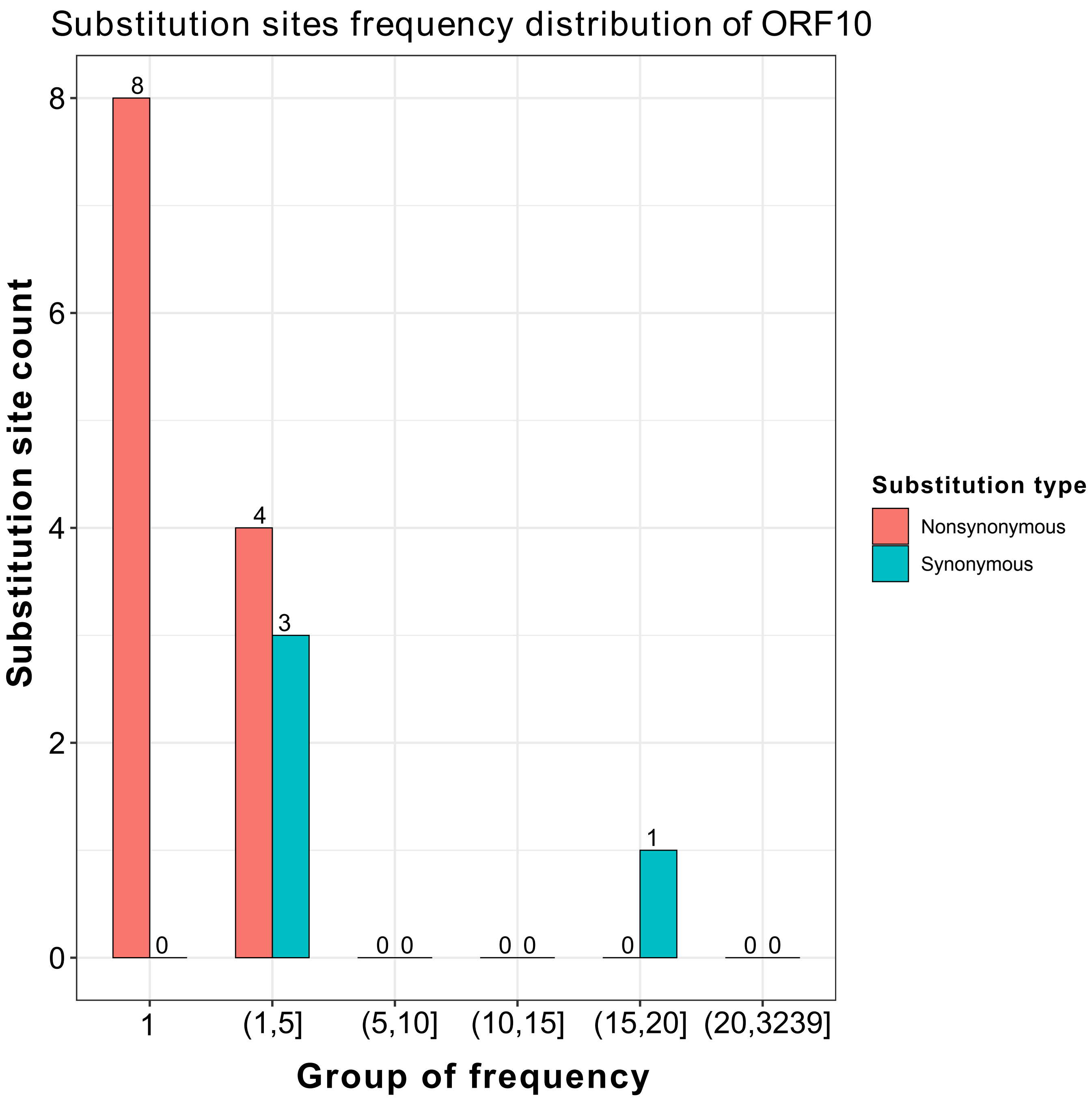

Substitution sites frequency distribution of ORF10
Substitution site count
Group of frequency
Substitution type
Nonsynonymous
Synonymous
8
4
3
1
0
0 0
0 0
0
0 0
0
2
4
6
8
1
(1,5]
(5,10]
(10,15]
(15,20]
(20,3239]
