## Supplementary figures and images for "Characterization of the substitution hotspots in SARS-CoV-2 genome using BioAider and detection of a SR-rich region in N protein providing further evidence of its animal origin"

### Supplemental Figure 2

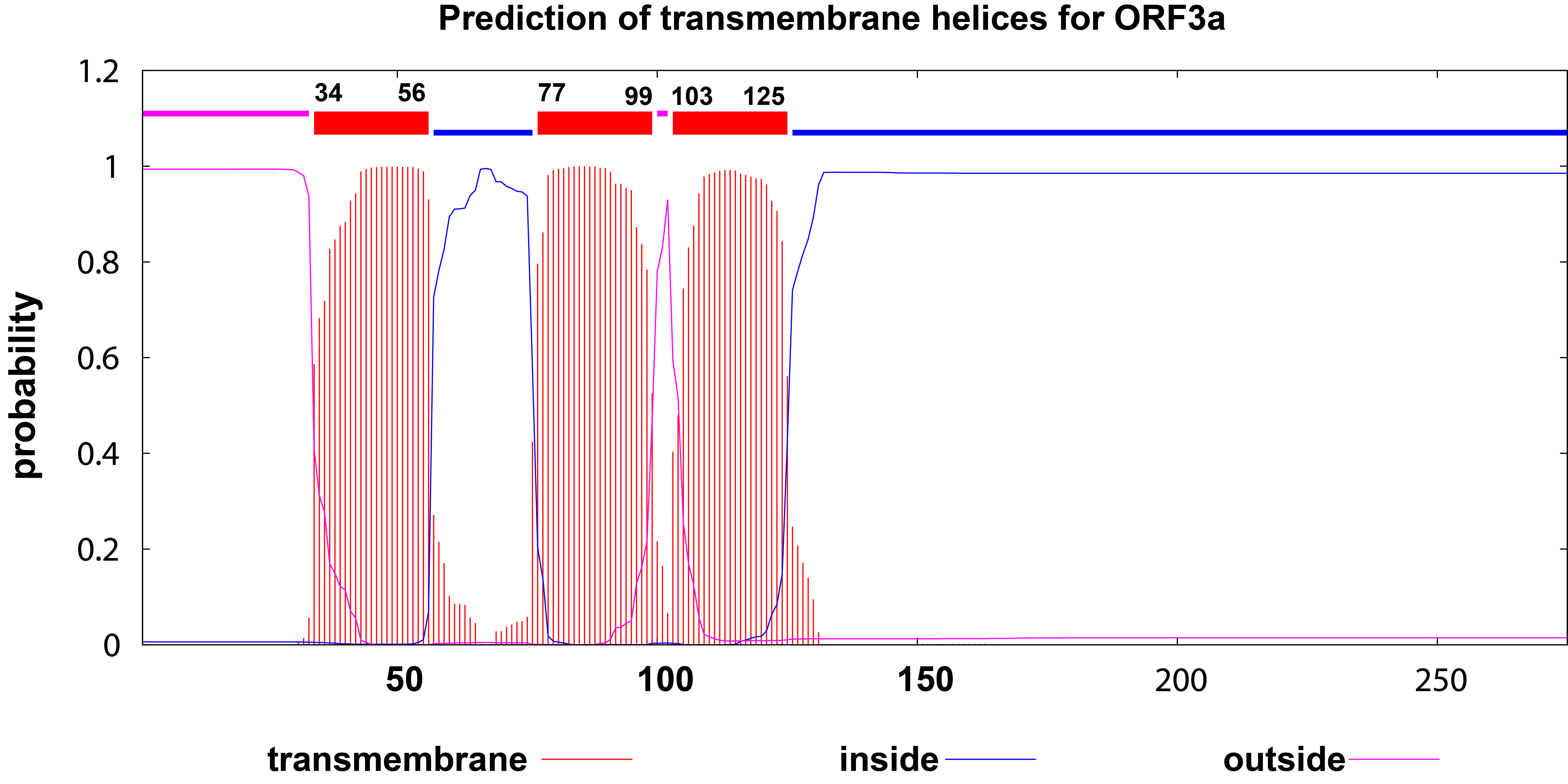

### Supplemental Figure 3

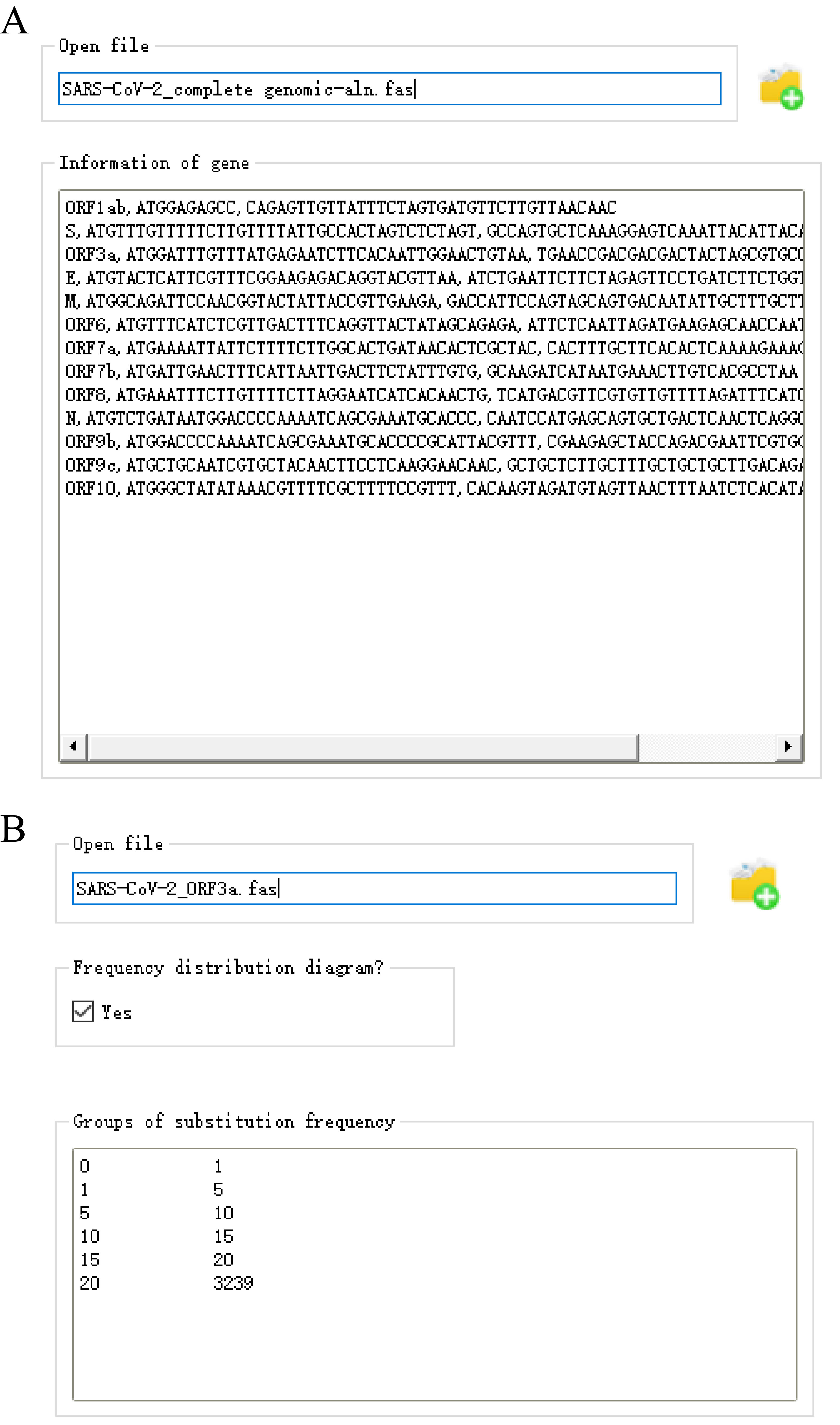
