## Supplemental material summary for "Characterization of the substitution hotspots in SARS-CoV-2 genome using BioAider and detection of a SR-rich region in N protein providing further evidence of its animal origin"

**Figure ledgend:**

**Supply Figure 1. Substitution frequency of synonymous or nonsynonymous sites in each gene encoded by SARS-CoV-2.** All the genes or ORFs of 1ab, S, ORF3a, E, M, ORF6, 7a, 7b, ORF8, N, ORF10 were analysed, respectively. The frequency of X axis indicates the number of variant strains at the substitution site, and the number one of first group represents that only one strain was mutated at this site. The Y axis represents the count of substitution sites corresponding to the range of substitution frequency. These frequency spectra were drawn using BioAider by specifying five different groups.

**Supply Figure 2. Prediction of transmembrane helices for ORF3a of SARS-CoV-2.** The sequence of ORF3a for prediction is EPI_ISL_402119.

**Supply Figure 3. Usage example of BioAider.** (A) Gene extraction for auxiliary sequence annotation. (B) Mutation analysis, note, the frequency distribution diagram of substitution sites is drawn based on the R program.
